# Systematic interrogation of mutation groupings reveals divergent downstream expression programs within key cancer genes

**DOI:** 10.1101/2020.06.02.128850

**Authors:** Michal R. Grzadkowski, Hannah Manning, Julia Somers, Emek Demir

**Affiliations:** Oregon Health & Science University, Portland, OR 97239

## Abstract

Genes implicated in tumorigenesis often exhibit diverse sets of genomic variants in the tumor cohorts within which they are frequently mutated. We sought to identify the downstream expression effects of these perturbations and to find whether or not this heterogeneity at the genomic level is reflected in a corresponding heterogeneity at the transcriptomic level. Applying a novel hierarchical framework for organizing the mutations present in a cohort along with machine learning pipelines trained on sample expression profiles we systematically interrogated the signatures associated with combinations of perturbations recurrent in cancer. This allowed us to catalogue the mutations with discernible downstream expression effects across a number of tumor cohorts as well as to uncover and characterize a multitude of cases where subsets of a genes mutations are clearly divergent in their function from the remaining mutations of the gene.

## INTRODUCTION

Each tumor faces a common set of obstacles arising from internal dynamics and external defense mechanisms^1^. Tumor cohorts, however, are replete with diverse yet recurrent tactics for overcoming these shared obstacles. Tumorigenesis can thus be perceived as a landscape within which each tumor navigates a unique, multidimensional path, weaving between segments trodden by other tumors. A number of the early breakthroughs in cancer treatment directly resulted from coarse demarcations of these paths into distinct subtypes based on “landmarks”—usually defined by mutations and/or markers derived from proteomic or transcriptomic data—that were then used to engineer subtype-specific treatments^2–4^.

Although these biomarker-based treatment matching criteria have proven effective in some precision medicine applications, there is a sizable subset of patients whose tumors harbor no discernible drug targets, thus diminishing their likelihood of successful treatment and survival^5–8^. Developing a more thorough understanding of the downstream effects of landmark events could therefore improve tailored treatment design outcomes. In particular, we envision a tactic which detects whether two genomic alterations (or combinations thereof) have a shared downstream effect, and can therefore be grouped together when weighing treatment options. This type of approach should also be able to detect whether two such alterations or groupings result in divergent transcriptional programs and can therefore be considered distinct. Despite recent efforts to profile the downstream effects of mutations recurrent in cancer, for most mutations we still know little about the programs they trigger. As a result, most clinical guidelines depend on only a limited subset of specific perturbations within a gene or on other coarse biomarker-based demarcations^9,10^. A clearer discernment of the convergences and divergences between the downstream programs present within cancer genes is thus a crucial prerequisite for addressing the challenges presently faced by precision oncology^11,12^.

Mutations of frequently altered genes often manifest as patterns of differential expression in other downstream genes. Such patterns are usually referred to as the transcriptomic signature or program associated with the mutation. It was previously shown that it is possible to generate transcriptomic signatures for common cancer drivers by training machine learning algorithms to predict which samples in a tumor cohort harbor their mutations^13–16^. A corollary to these results is that these mutation classifiers should also provide insight into the effects of the mutation in question. This hypothesis is supported by the correlations observed between these models’ predictions and other measurements of downstream activity including protein levels, response to drug treatment, and mutations in genes belonging to related cancer pathways^17,18^.

Further development of transcriptomic signatures is complicated by the dissimilitude of driver mutations within a gene^19,20^. Although genes such as BRAF carry one hotspot responsible for almost all mutations observed in the gene in tumor cohorts^21^, many genes implicated in tumor progression and proliferation have a widely distributed pattern of genomic alterations^22–25^. These mutations have varying degrees of impact and some are neutral. Moreover, it is not uncommon for different alterations within a gene to carry out diametrically opposite roles in cancer development depending on context^26^. In cases such as KRAS this property has already been exploited to engineer clinical interventions targeted to a specific KRAS hotspot rather than the gene as a whole^27^.

Can we measure these variable and divergent impacts? Consider the case in which a gene contains multiple groupings of mutations, each significantly divergent from the rest with respect to downstream impact. In this scenario, we would expect transcriptomic classifiers trained to predict the presence of mutations within individual groupings to be more accurate than a gene-wide classifier trained to predict the presence of any mutation of the gene. Conversely, if we do not observe increased classifier performance for subgroups, it is likely that they are convergent. Although subgroup-specific classifiers benefit to a certain degree from having a more uniform set of downstream effects to identify in a tumor cohort, they must also overcome the loss in statistical power inherent in characterizing a set of mutations present in a smaller proportion of available training samples. The discovery of mutation groupings robustly associated with better-performing classifiers within a gene would hence clearly present strong evidence of divergence.

The landscape of transcriptomic classifier accuracy across cancer genes’ mutation subgroupings should thus inform us about the convergent and divergent effects of these mutations. This understanding is useful for two immediate clinical purposes: estimating the likelihood that a variant of unknown significance has an effect similar to a previously characterized hotspot variant, and obtaining an informed grouping criteria for recurrent mutations to aid in the design of clinical trials and precision medicine guidelines. Based on these observations, we examined frequently altered genes in large tumor cohorts by systematically interrogating mutation subgroupings associated with improved classification performance. Instead of focusing on a single gene or pathway of interest, we sought to create a framework inspired by class-grouping approaches^28–30^ which would generalize well to the population of somatic alterations recurrent in cancer. Specifically, we test each gene for the existence of at least one good multinomial classifier by searching over a hierarchy of one-vs-rest binary classifiers. We confirm previous findings showing that it is possible to predict the presence of mutations associated with cancer using regression models trained on expression data and expand upon them to demonstrate the utility of taking into account the heterogeneity of mutation profiles within individual genes.

## MATERIALS AND METHODS

### Expression dataset preparation

We characterized the divergences present across alteration landscapes in cancer using transcriptomic signatures trained on a collection of publicly available tumor cohorts drawn from the METABRIC^31^, TCGA^32,33^, and Beat AML^34^ projects. In each of these cohorts, we filtered out samples for which either expression or mutation data was not available. Applying UMAP, a manifold-based unsupervised learning technique^35^, to the expression profiles of the remaining samples revealed clusters in cohorts such as TCGA-BRCA and TCGA-HNSC corresponding to molecular subtypes known to have unique transcriptomic profiles^36–38^ (Figure S1). To ensure that this heterogeneity at the molecular level did not confound our interrogation of heterogeneity at the genomic level, we partitioned cohorts containing subtypes associated with readily identifiable transcriptomic clusters. This yielded sub-cohorts such as METABRIC(LumA) and HNSC(HPV-) which were used alongside cohorts that did not require partitioning (Figure S2, Table S1).

### Enumeration of cancer gene mutation subgroupings in tumor cohorts

For clarity, we apply the term *point mutation* loosely to describe any genomic alteration involving a small number of nucleotides (*e.g.* SNPs, frameshifts, inframe insertions) while reserving the more general term *mutation* for the broader collection of perturbations spanning both point mutations and large-scale mutations such as copy number alterations (CNAs). We identified oncogenic and tumor suppressor genes based on their inclusion in the OncoKB database^39^. Our search for mutation groupings was restricted to the subset of these genes with point mutations present in at least 20 samples in any one of our cohorts. For each such gene, we enumerated subsets of its point mutations that could potentially have a biologically meaningful downstream transcriptomic signature.

Specifically, we hierarchically decomposed the variants of frequently mutated genes within a particular cohort. Each level in one of these hierarchies is associated with an attribute that can be used to characterize mutations in a non-overlapping, discrete manner, thus resulting in branches into which the mutations present in the cohort can be sorted. For example, one can build a simple two-level hierarchy ⟨Exon → Amino Acid Position⟩ that clusters a gene’s mutations according to genomic position by first grouping together mutations located on the same exon and then grouping together each exon’s mutations according to their amino acid position. Other attributes employed as hierarchical discriminators included overlap with a known binding domain, as well as the translation effect of the mutation (*e.g.* missense, nonsense, frameshift, etc.). Although our method can be applied to both point mutations and CNAs, here we focus on the former due to the richer set of attributes available for sorting them into our mutation trees.

For each cancer gene satisfying the 20-sample recurrence threshold in a cohort, we arranged its point mutations according to four hierarchies chosen to express useful biological priors about its mutations’ possible downstream effects. ⟨Exon → AA Position → AA Substitution⟩ is the two-level hierarchy described above that then further groups mutations at each amino acid position according to the specific amino acid that replaces the wild-type. This is useful for grouping together mutations that are located close to one another and can therefore be expected to bear a higher likelihood of having similar roles in downstream processes. Both ⟨SMART Domain → Form(base)⟩ and ⟨Pfam Domain → Form(base)⟩ first organize mutations according to their overlap with a known protein domain and then segregate mutations according to translational “form” (*i.e.* missense, nonsense, frameshift insertion, etc.) while grouping together insertions and deletions of forms such as frameshifts (hence “base”). These two trees enhance how the “closeness” of mutations is defined using a secondary source of information about structural units within the protein they affect and also incorporate information on the general nature of the perturbation caused by the mutation. Finally, ⟨Form → Exon⟩ considers unamalgamated mutation forms (*i.e.* frameshift insertions and deletions considered separately from one another) that are then grouped together by exonic position. This accounts for the possibility that, within particular genes, mutations of the same type are more likely to have similar downstream effects than mutations that are close to one another.

Each of these mutation trees can be used to generate a population of *mutation subgroupings*, defined as a branch or a combination of branches in a constructed hierarchy. With the mutation trees described above one can generate subgroupings such as

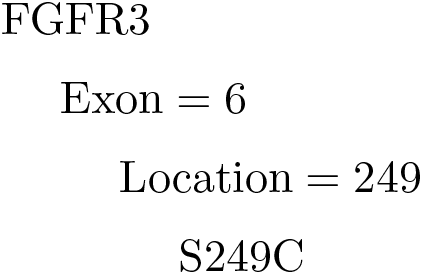

which consists of a single branch within the tree ⟨Exon → AA Position → AA Substitution⟩,

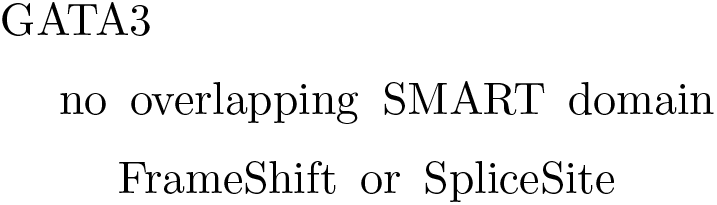

consisting of two branches of the tree ⟨Smart Domain → Form(base) → AA Substitution⟩, and

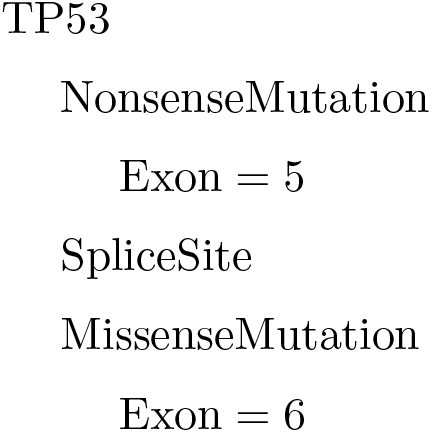

consisting of three branches in the tree ⟨Form → Exon⟩. While the first of these subgroupings simply represents the mutations belonging to a single hotspot of FGFR3, the other two represent less obvious groupings of the mutations of GATA3 and TP53 which we can interrogate for possible divergence from the remaining mutations of the gene.

Branches within subgroupings are combined using the union operation, *i.e.* all mutations on at least one of the branches are included in the subgrouping. To limit the number of mutation subgroupings to test, we only considered subgroupings consisting of at most two branches in one of the above hierarchies. Subgroupings were further filtered to only include those containing mutations present in at least 20 samples in a cohort, with each branch containing mutations present in at least 10 samples, and to remove subgroupings which included all the mutations of the corresponding gene.

Although these hierarchies allow for a fairly extensive search over the possible subsets of the mutations of a gene occurring in a cohort of samples, they do not offer a firm lower bound for finding the maximally divergent subgrouping. For our purposes, however, it is sufficient to detect at least one statistically significant divergent partitioning. Since we are systematically scanning all frequently altered genes across many cohorts, computational cost and statistical loss due to multiple hypothesis testing are limiting constraints. We found that our sampling heuristic based on biological priors can still elucidate multiple interpretable divergent subsets while pruning the search space down to a manageable size.

### Training and evaluating subgroup transcriptomic signatures

For each gene we trained a classifier to predict which samples in the cohort carried at least one point mutation on the gene—we refer to this as the gene-wide task. We then trained separate classifiers to predict which samples carried individual subgroupings’ mutations, referred to as the set of subgrouping tasks. Each task involved applying a logistic regression classifier utilizing the ridge regularization penalty to the given cohort’s expression data in order to generate binary labels for the samples that corresponded to whether they harbored a mutation in the subgrouping (see Supplementary Methods)^40^. To ensure that our subgrouping classifiers were identifying the downstream trans-regulatory effects of genomic perturbations and not solely their direct effects on the transcription of the corresponding gene and its genomic locality, we removed expression features associated with genes on the same chromosome as the gene whose mutations were being predicted. The output of each trained classifier was a continuous per-sample score denoting the classifier’s confidence that it was mutated in the subgrouping or gene, with higher values denoting greater confidence that a mutation was present. We measured a classifier’s ability to identify a transcriptomic signature for its assigned task using the area under the receiver operating characteristic curve metric (AUC) calculated using samples’ mean scores across ten iterations of four-fold cross-validation.

## RESULTS

### Subgrouping classifiers uncover alteration divergence in a breast cancer cohort

We found 38 genes with a total of 853 mutation subgroupings satisfying our classification task enumeration criteria using the 1017 METABRIC samples belonging to the luminal A sub-type of breast cancer. The transcriptomic signatures trained on these tasks revealed that many frequently mutated cancer genes within the METABRIC(LumA) cohort have readily identifiable downstream expression effects. Crucially, a sizeable subset of these genes contain subgroupings with expression signatures that diverge from those associated with the gene as a whole (Figure 1). For example, while it is easy to find a downstream effect in the expression data for GATA3 point mutations when they are considered as a whole (177 mutated samples; AUC=0.837), there are several subsets of GATA3 variants that produced even more accurate transcriptomic signatures: point mutations not assigned to an exon of GATA3 (in particular, splice variants) coupled with point mutations located on the 5th exon (79 samples; AUC=0.929), splice site mutations at codon 308 (43 samples; AUC=0.912), and frameshift mutations overlapping the zinc finger domain listed in the SMART database (36 samples; AUC=0.877).

**FIG. 1.**
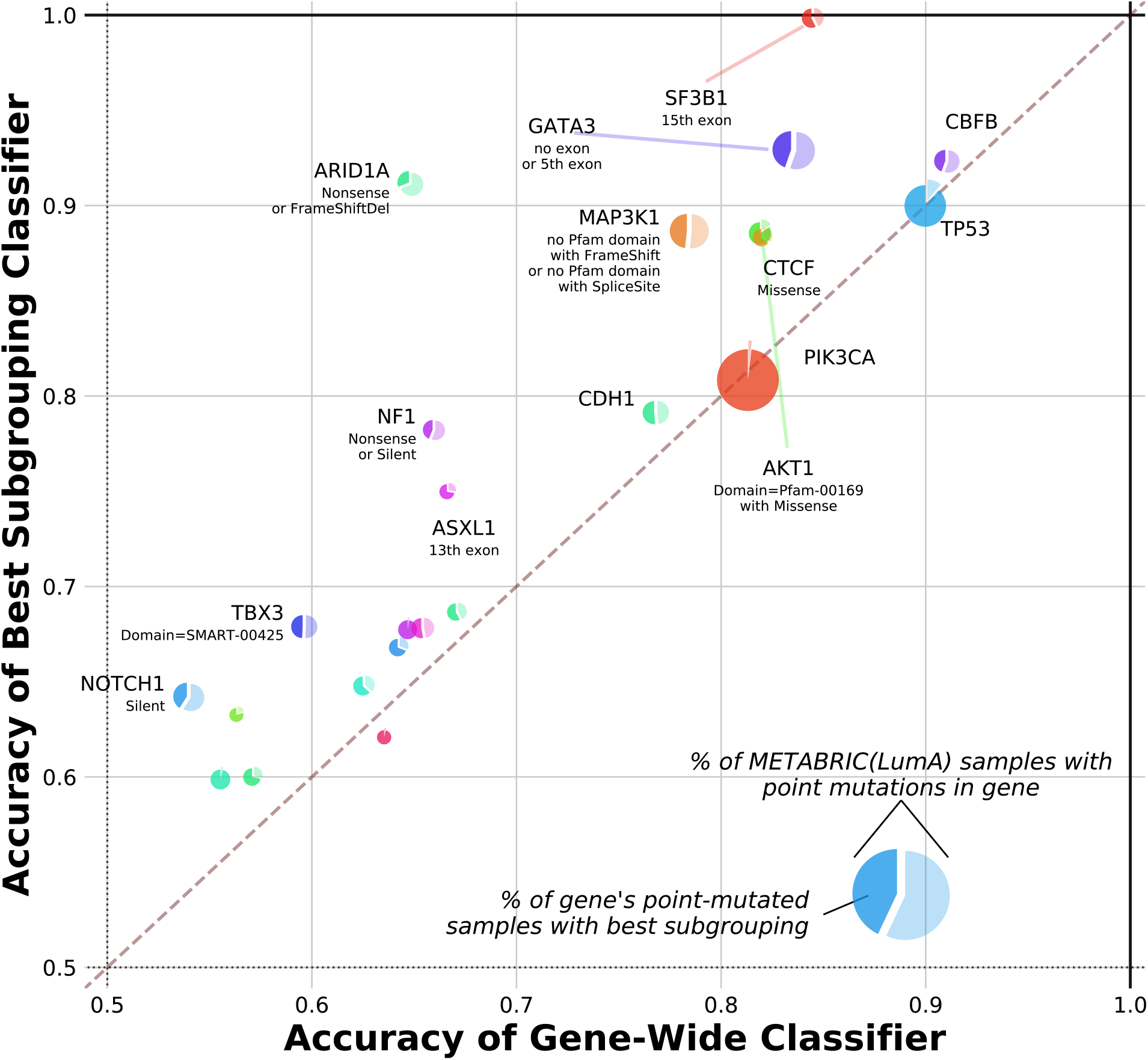
Divergent transcriptomic programs are a recurring feature of frequently mutated genes in breast cancer. 853 subgroupings within the point mutations of 38 genes with known links to cancer processes in METABRIC(LumA) were enumerated by grouping together variants with common properties. A logistic ridge regression classifier was trained to predict the presence of any point mutation in each of these genes as well as the presence of each enumerated subgrouping. Comparing the classification performance (AUC) for each gene-wide task (x-axis) to the best performance across all tested subgroupings of the gene (y-axis) reveals subgroupings within genes such as GATA3 and MAP3K1 with downstream effects that are consistently separable from the remaining mutations of the gene. The pie charts’ areas are proportional to the number of samples in the cohort that carry any point mutation of the corresponding gene; the darker slice inside each pie is scaled according to the proportion of these samples carrying a mutation in the best subgrouping, which for selected cases is described below the gene name.

These results are striking in that predicting the presence of a rarer type of mutation should, everything else being equal, be more difficult owing to decreased statistical power. Furthermore, while samples carrying any type of GATA3 mutation clearly have expression profiles distinct from those of samples that are wild-type for GATA3, our experiment demonstrates that it is also possible to find signatures that are able to consistently differentiate between different types of mutations within the GATA3 perturbational landscape. This is consistent with recent work showing that GATA3 mutations in breast cancer can be segregated according to their effect on the function of the GATA3 protein into subsets that broadly overlap with those identified as divergent above^41^.

Similar inferences can be made about the frameshift and splice site mutations of MAP3K1, which were found to segregate according to lack of overlap with the protein kinase domain (72 samples out of 149 mutated for MAP3K1; AUC=0.886 vs. AUC=0.786 for subgrouping task vs. gene-wide), as well as SF3B1 point mutations, within which mutations on the 15th exon were found to be highly divergent (27/47 SF3B1 mutants, AUC=0.999 vs. AUC=0.845). In the latter case, improved classification performance was primarily due to the presence of the K700E hotspot which accounted for all but one of these mutants, and could also be predicted nearly perfectly on its own (26/47 SF3B1 mutants, AUC=0.999). Moreover, the SF3B1 subgrouping which excluded silent mutations yielded a more modest boost in the quality of the transcriptomic signature (39/47 SF3B1 mutants, AUC=0.917), suggesting that SF3B1 variants can be ordered according to the strength of their downstream effects, with K700E mutations having the most significant impact. Several other genes, including CTCF and AKT1, were found to follow a similar pattern in which the best subgrouping was constructed by excluding silent mutations, using a single hotspot, or both.

The case of ARID1A was particularly noteworthy as classifiers struggled to find a signature when all of its mutations were considered together (68 samples; AUC=0.649), or even when restricted to predicting combinations of non-silent mutation types such as missense and nonsense (35 samples; AUC=0.550) or frameshift deletions and missense (34 samples; AUC=0.503). Only combining frameshift deletions and nonsense mutations of ARID1A into a subgrouping resulted in a major boost to classification performance for the gene (21 samples; AUC=0.911). This is consistent with the rarity of ARID1A missense mutations compared to nonsense mutations and frameshifts in other cancer types, suggesting that missense mutations are not selected for in general due to their lack of an effect on downstream processes^42,43^. This demonstrates our method’s ability to generate transcriptomic signatures for cancer genes which would be otherwise difficult to profile by identifying subsets of mutations that differ significantly in their behavior from other mutations on the same gene.

Intriguingly, we were unable to identify within-gene divergent subgroupings for TP53 and PIK3CA in METABRIC(LumA). Nevertheless, successfully training gene-wide classifiers for these well-known cancer drivers (AUC=0.901 and AUC=0.813 for the 221 TP53 and 488 PIK3CA point mutations respectively) lends further credence to our framework’s ability to identify down-stream effects where they would reasonably be expected to occur. It may be the case that nuanced yet consequential differences exist between the downstream expression effects of mutations within such genes, but that these differences are overshadowed by an expression program common to a sufficiently high proportion of the mutations. Similarly, whatever heterogeneity exists within the mutational profiles of these genes may be too granular to observe without access to larger tumor cohorts. For example, if TP53 variants could be rightly decomposed into dozens of distinct subgroupings, it would be difficult to find a transcriptional signature for each of these subgroupings individually within a cohort that only contains a total of 221 TP53-mutated samples. Our results are thus better interpreted as proving the divergence of some perturbational profiles rather than disproving the divergence of others.

### Subgrouping classifiers outperform random background benchmarks

Our approach to characterizing transcriptomic heterogeneity within the alteration profiles of cancer genes is based on testing as many mutation subgroupings as possible to identify those with divergent expression signatures. Although we have already demonstrated that this strategy can be gainfully applied to find such subgroupings, it is also clearly susceptible to multiple hypothesis testing—how can we be sure that the improvements in AUC we have observed are not simply the upper tail of the noise inherent in measuring the accuracy of a large population of classifiers? We thus devised several strategies to demonstrate the significance of our mutation classifier performances.

To establish a metric of confidence that the classification performance observed for the best subgroupings represented a significant improvement over using all point mutations for each gene and was not just a result of testing a multitude of subgrouping hypotheses, we derived a metric for comparing the AUCs of tasks to one another through down-sampling. A pool of 500 AUCs were generated for each task by randomly selecting 500 subsets of samples from the cohort and recalculating the AUC using solely the classifier scores returned for each of these sets of samples (see Supplementary Methods). This allowed us to interrogate the sensitivity of the AUCs we measured relative to the variation across the space of samples on which they were trained. Calculating the probability that a down-sampled AUC for each subgrouping task was higher than a down-sampled AUC for its associated gene-wide task trained on METABRIC(LumA) confirmed that our approach of considering subgroupings was particularly likely to yield a robust improvement in classification performance in the genes SF3B1 and ARID1A where all of the best found subgroupings’ down-sampled AUCs were higher than those of the gene-wide task (Figure S3). This down-sampling confidence metric also offered further support for the presence of divergence in genes such as GATA3 (conf=0.999), MAP3K1 (conf=0.999), and AKT1 (conf=0.967) when applied to the optimal subgrouping discovered in each case.

Furthermore, we cannot ascertain the significance of the AUCs we have observed in these classification tasks in isolation. Our prediction pipelines’ persistent ability to produce higher scores for samples in which a particular set of mutations is present leads us to claim that the set of mutations must have some biological relevance, but this relevance is difficult to establish without also comparing the classification performance against other sets of samples that could have been selected from the cohort to construct classification tasks. We thus created a set of classification tasks to predict the presence of randomly chosen sets of samples of the same size as the mutation subgroupings we previously tested. The distribution of AUCs for this null background set of tasks was markedly lower than the corresponding distribution for tasks related to cancer genes, confirming that the mutation labels associated with cancer genes encode a significant amount of information relative to randomly-chosen labels (Figures S4A-B). For each gene we also created a gene-specific null background set of classification tasks by randomly selecting subsets from the collection of samples carrying any point mutation of the gene. The performance observed for these tasks revealed that in cases such as GATA3, AKT1, and FOXA1 our hierarchical organization of the mutations in each gene yielded better subgroupings than those that could be found by simply picking subsets of mutations occurring on the gene at random as measured by the down-sampling confidence metric discussed above (Figure S4C). This underlines the utility of leveraging the various attributes of mutations as a biological prior for clustering them together into subgroupings with more uniform downstream transcriptomic effects.

### Mutation prediction performance is robust with respect to choice of classification algorithm

Since our method requires scanning a sizeable population of subgroupings, we opted to use a linear ridge regression classifier that efficiently scales up to a large number of tasks. However, this choice of algorithm can potentially prevent us from detecting nonlinear transcriptomic signatures. Our tuning regime was also designed to be fairly straightforward in order to reduce computational load, testing only eight ridge regularization hyper-parameter values. To find whether our mutation prediction results in METABRIC(LumA) were affected by these efforts to reduce the computational cost of our classification pipelines, we repeated the above experiment with radial basis function support vector^44^ and random forest classifiers^45^ as well as with a larger tuning grid for the ridge regression classifier.

These more complex classifiers failed to produce improved AUC performance across our prediction tasks and did not affect the efficacy of subgrouping tasks relative to gene-wide tasks despite taking up to an order of magnitude longer to run to completion for the cohort sample sizes considered (Figure S5). The Spearman correlation of AUCs across non-random classification tasks was 0.941, 0.893, and 0.984 for the support vector, random forest, and large-tuning-grid classifiers respectively against the AUCs measured using the original linear regression approach, further demonstrating our results’ invariance to the machine learning algorithm used. The mutational profile heterogeneity we observe is thus not a by-product of the behavior specific to any one particular classification method, and can be observed using a relatively simple learning framework. That linear regression is sufficient for successful prediction in this context is likely due to the relatively small size of the available tumor cohorts. Even if more complex relationships between genomic perturbations and expression levels are indeed present in breast cancer, it is likely difficult to characterize them without more statistical power than is available in a population of only 1017 samples.

### Enlarging the subgrouping search space does not significantly alter relative classifier performance

Relaxing the parameters of our subgrouping enumeration heuristic to allow for a larger search space of 7598 subgroupings that included those composed of up to three branches of at least five samples did not uncover a significant number of cases of divergent subgroupings that had not already been found using the original criteria (Figure S6). In cases such as AKT1 and SF3B1 the best possible subgrouping had clearly already been identified due to the limited number of combinations of mutation groupings we could test given the small number of samples carrying any of their point mutations. MAP3K1 and PIK3CA exhibited very modest improvements in classification performance using the enlarged pool of subgroupings (AUCs of 0.895 vs. 0.886 and 0.817 vs. 0.813 respectively for expanded vs. original subgrouping search spaces) which were not sufficient to justify a ninefold increase in computational cost. Nevertheless, the fact that the best subgrouping of TP53 from this larger set converges even closer to the gene-wide set of mutations lends greater credence to the difficulty of finding a divergent set of mutations within this gene. There were some genes, including CDH1 and PTEN, that benefited from our deeper subgrouping search. This is likely due to the fact that in both of these genes the mutations are well-spread, with no single hotspot accounting for more than 10% of the mutations occurring within them in METABRIC(LumA).

We also integrated copy number alterations (CNAs) in our subgroupings. For every enumerated subgrouping, we created up to two additional classification tasks using the point mutations in the subgrouping combined with deep deletions or deep amplifications of the same gene in cases where one or both of these types of CNAs were present in at least five cohort samples. However, this did not improve classification performance for cases such as GATA3, SF3B1, and AKT1 within which divergent subgroupings had already been found when not including CNAs in the set of mutations to predict (Figure S7). In genes such as TP53 and MAP3K1 there were simply not enough deep CNAs present in the METABRIC(LumA) cohort for us to test any subgroupings which included them. On the other hand, we found that using deep deletions along with missense mutations on the catalytic domain and frameshifts on the C2 domain of PTEN led to a well-performing transcriptomic signature (38 samples; AUC=0.762) compared to both using all PTEN point mutations (51 samples; AUC=0.655) or using all PTEN point mutations and deep deletions (67 samples; AUC=0.744). Including CNAs in our subgrouping enumeration also allowed us to better characterize genes such as ERBB2 (HER2) where using deep amplifications on their own produced a superior expression signature (43 samples; AUC=0.820) relative to using all point mutations of the gene (38 samples; AUC=0.671) or all point mutations in conjunction with deep gains (80 samples; AUC=0.722).

### Breast cancer cohorts exhibit concordant divergence characteristics

We performed the same analyses on TCGA-BRCA data to test whether the mutation grouping behavior in METABRIC generalizes to other breast cancer cohorts and is not simply an artefact of expression patterns specific to METABRIC or of over-fitting within our classification tasks. The subgrouping enumeration procedure described above was repeated with the TGCA-BRCA luminal A sub-cohort consisting of 499 samples to identify 16 cancer genes containing 238 subgroupings of which 14 genes and 222 subgroupings had also been enumerated in METABRIC(LumA). Training and evaluating classification tasks predicting the presence of these subgroupings using TCGA-BRCA(LumA) expression data revealed transcriptomic signature characteristics broadly concordant with what was observed in METABRIC(LumA), with a Spearman correlation of 0.734 across AUCs recorded for non-random subgroupings enumerated in both cohorts (Figure 2). This is despite the fact that the expression calls in the TCGA-BRCA(LumA) cohort were made in an independent setting using next-generation sequencing profiling as opposed to microarrays.

**FIG. 2.**
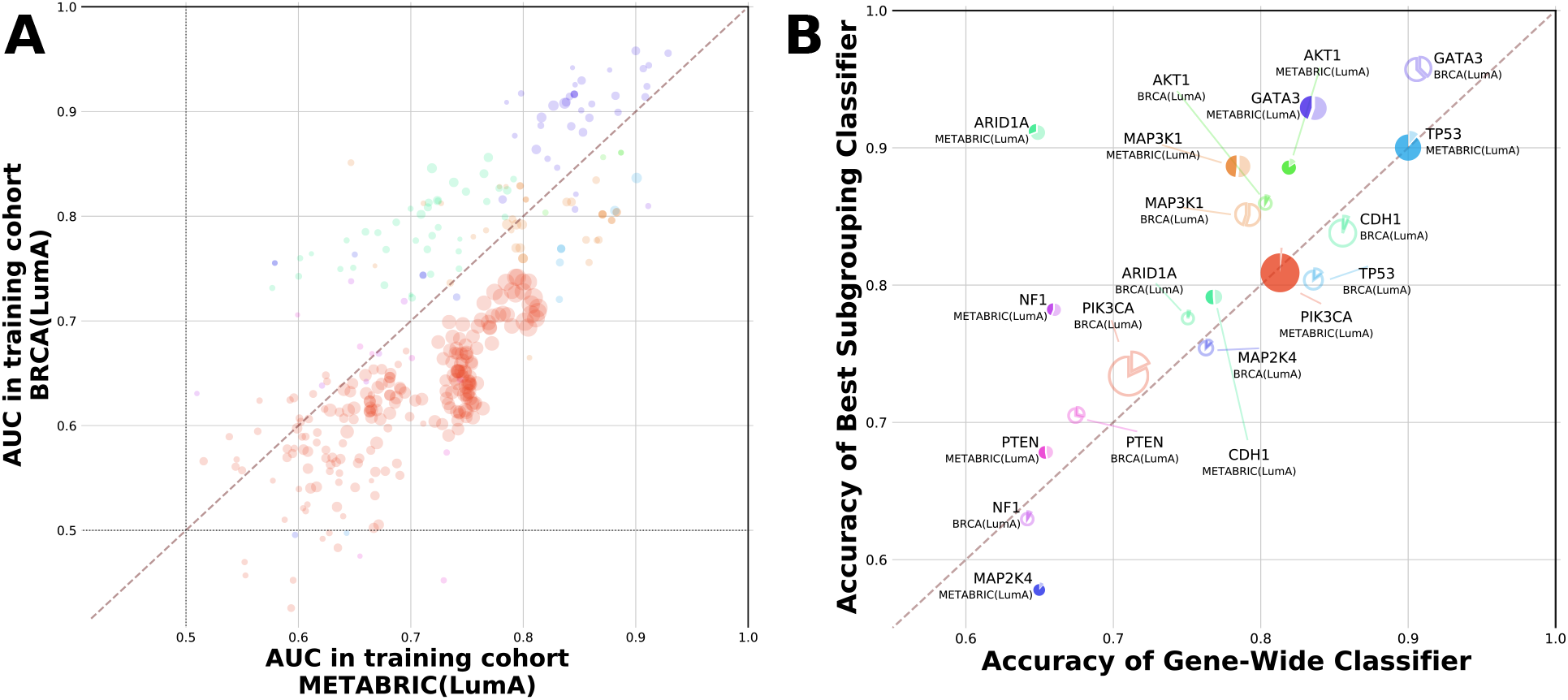
Subgrouping performance is consistent across breast cancer cohorts. Cancer gene subgrouping enumeration and classification was repeated using the luminal A sub-cohort of TCGA-BRCA. The colors for genes’ plotted points and pie charts correspond to those in Figure 1. (**A**) Prediction AUCs for gene-wide classification tasks and subgrouping tasks enumerated in both METABRIC(LumA) (x-axis) and TCGA-BRCA(LumA) (y-axis). Larger point size indicates a higher joint proportion of mutated samples (calculated as the geometric mean of the two cohort proportions). (**B**) Comparison of relative subgrouping performance (AUC) between cancer genes profiled in TCGA-BRCA(LumA) (filled-in pie charts) versus those profiled in METABRIC(LumA) (hollow pie charts).

In particular, genes such as GATA3, MAP3K1, and AKT1 which contain divergent subgroupings in METABRIC(LumA) exhibit the same behavior in TCGA-BRCA(LumA), while genes like TP53, PIK3CA, and CDH1 without such subgroupings in METABRIC(LumA) also lack them in the counterpart TCGA cohort. Furthermore, subgroupings found to be divergent in one cohort tended to also be identified as divergent in the other cohort (Figure S8). The set of missense mutations overlapping with the Pleckstrin homology domain was found to have the best AUC across AKT1 subgroupings in both METABRIC(LumA) and TCGA-BRCA(LumA), while the best subgroupings of GATA3 in each cohort also perform much better than the gene-wide task in the other cohort. An outlier in this regard was MAP3K1, in which the set of missense and frameshift mutations was found to have a strong relative AUC in METABRIC(LumA) (AUC=0.883 vs. AUC=0.786 for the gene-wide task), but did not exhibit similar performance in TCGA-BRCA(LumA) (AUC=0.804 vs. AUC=0.792 for the gene-wide task). A similar discordance was also observed in the opposite direction where the best MAP3K1 subgrouping identified in TCGA-BRCA(LumA) performed poorly in METABRIC(LumA) (AUC=0.851 vs. AUC=0.647 for the subgrouping of missense and nonsense mutations). Nevertheless, the discovery of at least some divergent subgroupings of MAP3K1 in each cohort suggests a consistent pattern of heterogeneity within its alterations, and that the set of mutations driving this divergence is not well-characterized by the particular combinations of mutation annotation levels over which we chose to enumerate subgroupings. ARID1A, notable as the case where divergent behavior was found in one breast cancer cohort but not the other, can be explained by the fact that the incidence of ARID1A mutations varies significantly between the two cohorts: in METABRIC(LumA) 6.7% of samples carry a point mutation in the gene, while in TCGA-BRCA(LumA) only 4.2% do. Since there are only 21 total mutated samples in the latter case, none of the ARID1A subgroupings that our method enumerated in METABRIC(LumA) satisfied the sample frequency threshold in the TCGA-BRCA(LumA) sub-cohort.

To further validate the generalizability of subgrouping performance, we applied the models trained to predict subgroupings found in METABRIC(LumA) to the TCGA-BRCA(LumA) cohort and vice versa. We found that the AUCs for these classifiers in the previously unseen cohort were similar to the cohort on which they were trained (Figure S9), with a Spearman correlation of 0.848 between the original AUCs for models transferred from METABRIC(LumA) to TCGA-BRCA(LumA) and their AUCs in the transfer context, and a Spearman correlation of 0.823 between the original and transfer AUCs for models trained on TCGA-BRCA(LumA) and transferred to METABRIC(LumA). Furthermore, for genes such as GATA3 and AKT1 with divergent subgroupings in the two breast cancer cohorts, these subgroupings also outperformed the corresponding gene-wide classifier in this transfer setting (Figure S10).

The robustness of our findings was further underlined by obtaining comparable results when running the same experiment using TCGA-BRCA(LumA) expression calls produced by kallisto^46^ as input rather than calls produced by RSEM^47^. Similar results were also obtained when using other combinations of the subtypes present in breast cancer instead of solely luminal A in both TCGA-BRCA and METABRIC (Figure S11). We thus conclude that the advantages of considering subgroupings within genes to model downstream transcriptomic effects are persistent when exposing these models to as yet unseen datasets, and that these mutation models generalize well across different breast cancer cohorts and expression quantification methods.

### Divergent cancer gene subgroupings are present across a variety of cancer types

To further interrogate the presence of divergent alteration profiles across different tumor contexts, we repeated our enumeration and classification steps across the fourteen other cohorts in TCGA with a sufficient number of samples as well as the Beat AML cohort. In total, 6530 subgroupings across 160 different genes were tested using the TCGA cohorts, in addition to the 853 subgroupings across 38 genes tested in METABRIC(LumA) and the 132 subgroupings across 14 genes tested in Beat AML. This revealed that gene-wide expression signatures can be trained for a number of cancer genes in most oncological contexts (Figure 3). Furthermore, divergent subgroupings are a feature of not just breast cancer but of many other tumor types as well (Figure S12). A multitude of genes exhibit at least one subset of point mutations with a robust transcriptomic signature that significantly outperforms the gene-wide signature (Table I).

**FIG. 3.**
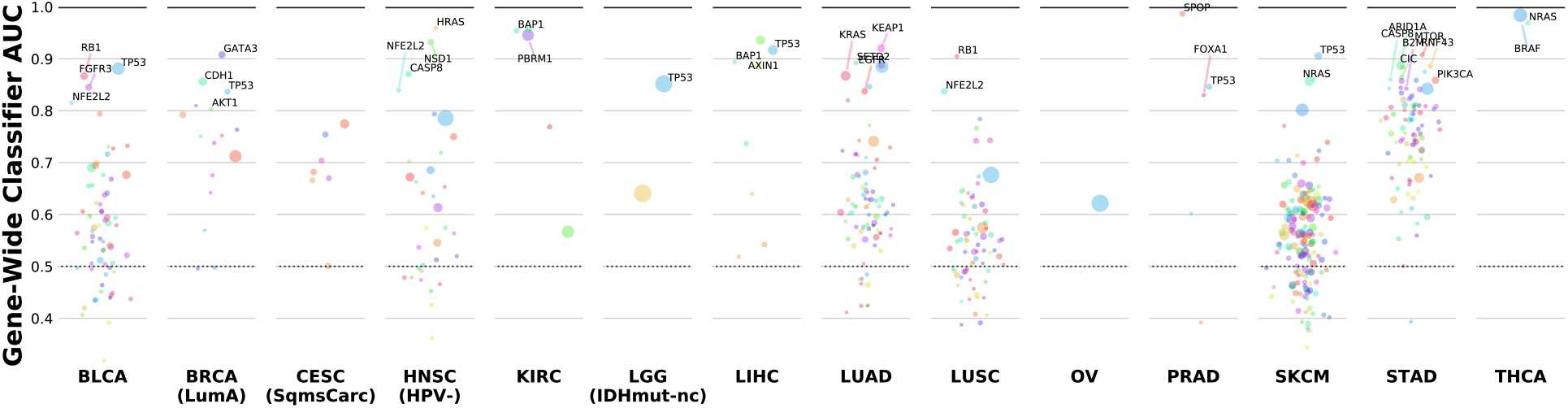
Many cancer genes’ point mutations have identifiable expression signatures. Our experiment attempted to predict the point mutations of a total of 192 cancer genes across 15 TCGA tumor cohorts using transcriptomic profiles. Shown are the AUCs for all 555 of these gene-wide tasks, with particularly well-performing classifiers highlighted. Point size corresponds to number of point-mutated samples.

**TABLE I.** Cataloguing cancer genes with divergent subgroupings across TCGA cohorts. The subgrouping enumeration and classification tasks were applied to each TCGA cohort meeting our selection criteria. Notable cases of genes containing subgroupings with significantly better AUCs than the corresponding gene-wide task are listed above.

| Cohort | Gene | Mutation Subgrouping | AUCs<sup>†</sup> | Samps<sup>‡</sup> |
| --- | --- | --- | --- | --- |
| BLCA | FGFR3 | any mutation on 9th exon or S249C | 0.919 vs. 0.845 | 49/61 |
|  | NFE2L2 | 2nd exon | 0.939 vs. 0.815 | 20/27 |
| HNSC(HPV-) | FBXW7 | WD40 repeat domain | 0.887 vs. 0.794 | 26/33 |
|  | NSD1 | frameshift w/o domain or nonsense w/o domain | 0.976 vs. 0.932 | 32/52 |
| LIHC | CTNNB1 | 3rd exon | 0.968 vs. 0.936 | 78/94 |
| LUAD | EGFR | inframe del on any exon or missense on 21st exon | 0.934 vs. 0.837 | 51/73 |
| LUSC | NFE2L2 | 2nd exon | 0.896 vs. 0.837 | 67/75 |
| PRAD | FOXA1 | fork head domain | 0.957 vs. 0.830 | 22/31 |
| STAD | APC | frameshift w/o domain or nonsense w/o domain | 0.854 vs. 0.756 | 21/53 |
|  | ATM | frameshift w/o domain or missense w/o domain | 0.882 vs. 0.782 | 31/46 |
|  | CREBBP | any mutation w/o acetylation domain | 0.898 vs. 0.821 | 25/44 |
|  | PIK3CA | missense on C2 domain or w/o domain | 0.900 vs. 0.859 | 40/70 |
|  | SPEN | frameshift w/o domain or silent w/o domain | 0.868 vs. 0.811 | 29/51 |

<sup>†</sup> given as (AUC of listed subgrouping) vs. (AUC of gene-wide classifier)
<sup>‡</sup> number of samples in the subgrouping task vs. in the gene-wide task

For example, within the context of prostate adenocarcinoma (TCGA-PRAD), mutations of FOXA1 that overlap with the fork head binding domain are much easier to predict than all FOXA1 point mutations taken together (22/31 FOXA1 mutants; AUC=0.957 vs. AUC=0.830; conf=0.988). This is consistent with the importance of such domains in guiding the regulatory functions of transcription factors as well as with previous characterizations of the functional divergences present within FOXA1 mutations in prostate cancer cohorts^48–50^. In the HPV-subtype of head and neck squamous carcinomas (TCGA-HNSC), NSD1 was found to have a divergent subgrouping consisting of frameshifts and nonsense mutations not overlapping with a protein domain (32/52 samples, AUC=0.976 vs. AUC=0.932, conf=0.970). Thus we can deduce that the 18 missense mutations of NSD1 present in the cohort have a much weaker downstream effect, especially since the same subgrouping with nonsense mutations replaced by missense mutations had poor classification performance (24/52 samples, AUC=0.818). EGFR contains two major hotspots (E746-A750del and L858R) in the lung adenocarcinoma cohort (TCGA-LUAD) which form the bulk of the samples in the best found subgrouping (51/73 samples, AUC=0.934 vs. AUC=0.837, conf=0.995). This implies that these two loci have similar or at the very least highly complementary impacts on the transcriptome.

We also found that genes frequently mutated in multiple cohorts tended to be consistent in the overall structure of their alterations’ downstream effects. For instance, we can produce a well-performing signature for TP53 variants in multiple cancer cohorts including melanoma (67 mutated samples in TCGA-SKCM; AUC=0.905), bladder cancer (200 muts in TCGA-BLCA; AUC=0.881), and lung adenocarcinoma (264 muts in TCGA-LUAD; AUC=0.885) in addition to the signature found in luminal A breast cancer already described above. However, TP53 mutations do not exhibit divergence in any of these cancers, as no subgrouping’s transcriptomic signature was found to be significantly divergent from that of TP53 mutations as a whole (down-sampled confidence scores of 0.66, 0.04, and 0.04 respectively) (Figure S13A). Likewise, PIK3CA has no detectable divergence in 7 out of 9 cohorts where it satisfied the mutation recurrence threshold and only a very weak divergence in stomach and small-cell lung cancer (Figure S13B).

In contrast to this, NFE2L2 is associated with both a robust downstream signature and significant divergence in all three cohorts in which its subgroupings were enumerated (TGCA-BLCA: 20/27 muts, AUC=0.939, conf=0.97; TCGA-LUSC: 67/75 muts, AUC=0.896, conf=0.97; TCGA-HNSC(HPV-): 21/31 muts, AUC=0.898, conf=0.84) (Figure S13C). Meanwhile, cases where strong divergence was observed in some cohorts but not others tended to be driven by varying mutation frequencies and types rather than inconsistent patterns of downstream effects. For example, the divergent subgrouping of FOXA1 in TCGA-PRAD described above has only 24 samples bearing point mutations in TCGA-BRCA(LumA) and none in METABRIC(LumA) (Figure S13D). The cause of this discrepancy remains beyond the scope of this paper, but we speculate that it is primarily contingent upon differences in sequencing platforms and pipelines.

The transfer validation method that was used to compare the performance of trained models between our two breast cancer cohorts was extended across all of the cohorts we used for training. Transferring trained classifiers across disease contexts revealed that transcriptomic models for cancer genes such as TP53 generally perform well even when they are applied to a tumor type different from that in which they have been trained (Figure S14). This reflects both the robustness of our classification pipelines and the ubiquitous nature of the downstream effects associated with TP53 perturbations. On the other hand, even high quality PIK3CA models do not migrate as well among tumor types, suggesting that recurrent PIK3CA mutations may result in unique downstream transcriptional signals predicated on the unique cancer context in which they developed. In NFE2L2, subgrouping models performed well in transfer validation and outperformed models trained using all of the gene’s point mutations in transfer contexts (Figure S15). These subgrouping models are therefore especially likely to preserve their performance when applied to novel cohorts or patient samples, which is especially important in a variety of clinical settings where they would be implemented.

### Subgroupings outperform mutation subsets chosen using variant significance metrics

We have already compared the classification performance with our mutation subgroupings against the performance when using all point mutations for the corresponding gene, as well as against the performance when using sets of samples chosen at random from both the training cohort as a whole and the set of samples carrying any point mutation on the gene. To further validate our approach, we compared the performance of classifiers tasked with predicting the presence of our mutation subgroupings against those predicting subsets of mutations constructed using existing metrics designed to capture the impact of mutations on cancer processes. For each gene with enumerated subgroupings in a cohort we thus created a classification task for each possible threshold value of the PolyPhen and SIFT scores^51,52^ assigned to its variants that resulted in a unique set of at least 20 samples carrying a mutation of the gene satisfying the threshold. This allowed us to evaluate the relative efficacy of the transcriptomic signature trained using a subgrouping containing *n* mutated samples of a gene against that of a signature trained using a subgrouping containing the top *n* mutants according to PolyPhen or SIFT wherever these scores were available.

We found that in cases such as EGFR in TCGA-LUAD and NFE2L2 in TCGA-HNSC our method of discovering subsets of mutations outperformed any possible choice of cutoff of the above metrics (Figures 4A-D). For the former, the aforementioned subgrouping (composed chiefly of the E746-A750del and L858R hotspots) exhibited significantly better performance than the best found PolyPhen cutoff of ≥ 0.957 which included the top 34 EGFR-mutated samples in TCGA-LUAD according to this metric (AUC=0.934 vs. AUC=0.862; conf=0.974), as well as the best found SIFT cutoff of ≤ 0.12 which included 37 EGFR mutants (AUC=0.867; conf=0.968). This is explained by the fact that the E746-A750del mutation is not assigned a score by either PolyPhen or SIFT, and thus is not included in any of the corresponding threshold-based classification tasks, hence exposing the limitations of these metrics relative to our more flexible method of gauging the downstream effects of mutations. On the other hand, in the case of NFE2L2 in TCGA-HNSC, the difference between our best subgrouping consisting of the 21 NFE2L2 mutants with perturbations on the gene’s second exon and the best found cutoff for PolyPhen (≥ 0.999) was due to a mutation present in a single sample (V32G) scored as benign by PolyPhen, while the difference between this subgrouping and the best found cutoff for SIFT (≤ 0.05) was largely due to three mutants on other exons (D457G, L562F, D570N) that were also predicted as being deleterious according to SIFT. Despite these small differences in the composition of the classification tasks, the improvement in prediction accuracy was considerable in both cases when using our subgrouping (AUC=0.898) versus these metrics (PolyPhen AUC=0.848; conf=0.757, SIFT AUC=0.841; conf=0.657). From these findings we conclude that the downstream expression effects of mutations on the second exon of NFE2L2 are likely to be uniform with respect to one another but divergent from other deleterious mutations of NFE2L2.

**FIG. 4.**
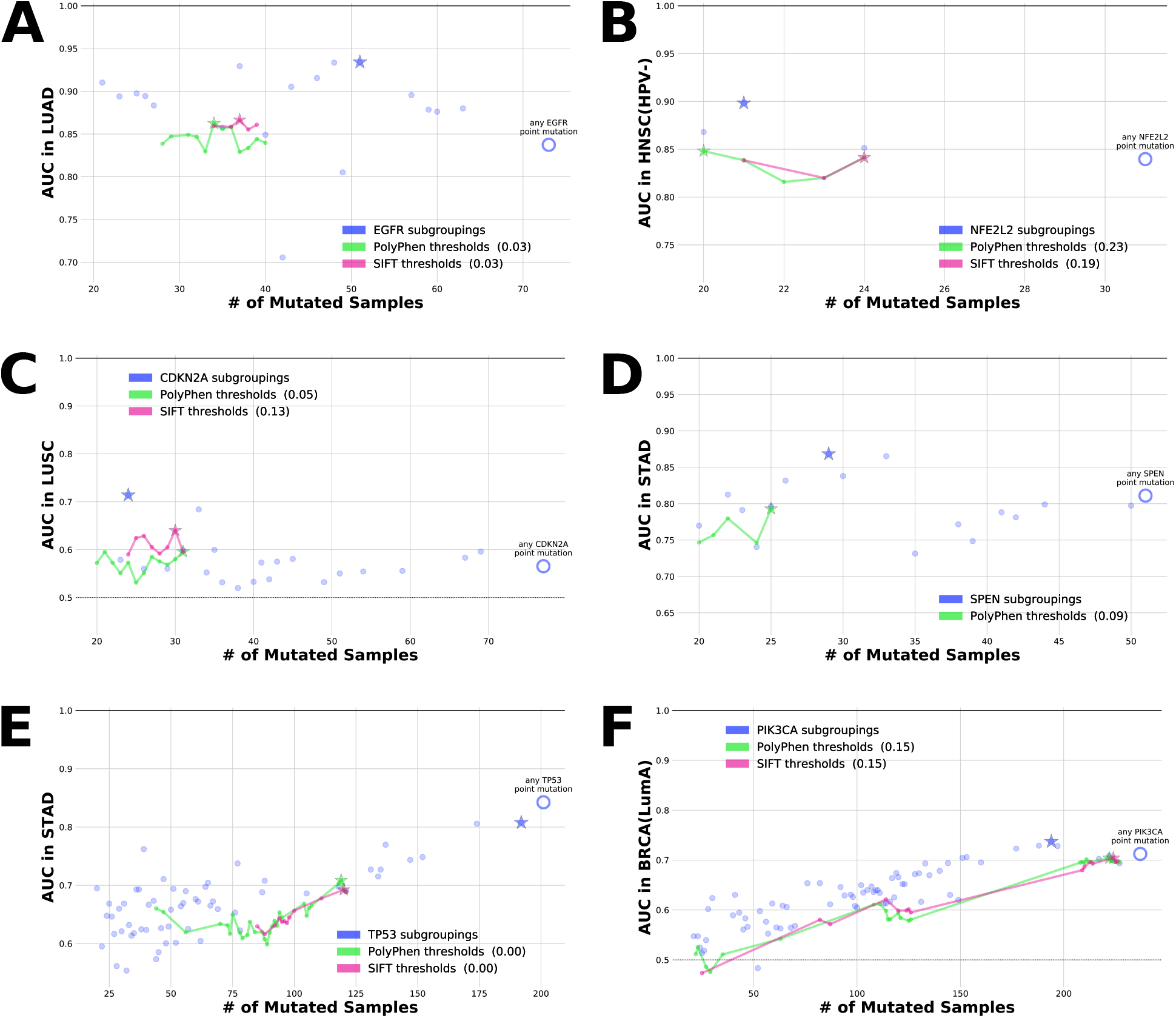
Comparing subgroupings against mutation subsets defined by other tools for measuring variant significance. Classification tasks were created in which the top *n* samples according to the value of various continuous mutation properties were treated as a discrete subgrouping. Using the same training and testing regime as before, we compare the AUCs for these tasks to those for subgrouping tasks created using our original discrete approach. This revealed cases where using subgroupings was clearly superior to using these metric cutoffs (**A**-**D**), as well as cases where neither subgroupings nor cutoffs significantly outperform the genewide classifier (**E**and **F**). Legend labels are annotated with the sub-sampled confidence score of the best subgrouping (star marker) against the best set of mutations chosen using the given cutoff.

Using PolyPhen and SIFT cutoffs also failed to find divergence within genes such as TP53 and PIK3CA where we had not discovered any divergent subgroupings (Figures 4E-F). This lends further credence to the possibility that the divergences within the mutation profiles of these genes, if they do exist, are overshadowed by a common expression program. When we performed this comparison for all cases with divergent subgroupings, we found that many of these subsets could not have been identified using either PolyPhen or SIFT (Figure S16). Our subgrouping enumeration method can thus outperform other approaches for evaluating the potential impact of oncogenic mutations, and highlights the importance of incorporating a variety of biological priors when characterizing the relationships between genomic perturbations and their tumorigenic effects.

### Subgrouping classifier output reveals the structure of downstream effects within cancer genes

Comparing the performances of transcriptomic signatures for different subsets of mutations within cancer genes has allowed us to identify divergences within them. However, this analysis does not on its own pinpoint the nature of the differences within these genes’ mutations that are responsible for this observed heterogeneity—a subgrouping could have a transcriptomic profile that diverges from that of its parent gene for a variety of reasons. For instance, it is possible that the mutations of the gene not belonging to the subgrouping are functionally silent. Another possibility is the existence of multiple transcriptomic programs within the gene that are complementary or orthogonal to one another, each of which can be uniquely mapped to a subset of the gene’s mutations. We thus investigated the output of the signatures we trained for these subgroupings to better understand the mechanisms driving the downstream transcriptomic effects of tumorigenic alterations.

For selected subgroupings that had been identified as divergent in the cohorts we included in our experiment we examined the mutation scores their expression classifiers returned for the mutations on the same gene not belonging to the subgrouping (Figure S17). This helped us to characterize the relationships that were responsible for the observed divergence between each subgrouping and the remaining mutations on the gene in which they were found. For example, we were able to confirm that the mutations falling outside of the best found subgrouping of missense mutations overlapping the Pleckstrin homology domain within AKT1 in METABRIC(LumA) behave like wild-type samples according to our classifiers’ scores, which is consistent with the fact that most of these are synonymous substitutions.

More surprising was finding that ARID1A missense mutations in METABRIC(LumA) are predicted by both the best found subgrouping’s classifier and the gene-wide classifier as behaving like ARID1A wild-types and synonymous mutations (Figure S18A). This is strongly suggestive of the neutrality of these mutations with regards to downstream transcriptomic processes, especially relative to the nonsense and frameshift mutations our framework identified as constituting a divergent subgrouping. Furthermore, treating ARID1A missense mutations as equivalent to ARID1A wild-types and synonymous substitutions resulted in a classifier that was better able to separate samples with active ARID1A mutations from the remainder of the cohort, with far fewer nonsense and frameshift ARID1A mutants being assigned low scores by the optimal subgrouping’s classifier compared to the gene-wide classifier. Similar behavior was observed for mutations on exons other than the second within NFE2L2 in TCGA-BLCA and TCGA-LUSC as well as mutations not belonging to the L858R or E746-A750del hotspots within EGFR in TCGA-LUAD (Figures S18B-D). In all three of these cases our method of organizing subsets of mutations within a gene allowed us to differentiate between mutations with and without a downstream effect, which explains how we were able to train a mutation classifier with a higher accuracy than the gene-wide model.

In other genes, our classification framework uncovered a blurrier distinction between active and inactive mutations. SF3B1 mutants not belonging to the K700E hotspot which constituted the bulk of the best found subgrouping for the gene in METABRIC(LumA) were assigned scores between that of subgrouping and wild-type samples. Further investigation revealed that samples with SF3B1 mutations on the 14th exon closest to the K700E hotspot were especially likely to be seen as having higher scores than mutations on the other exons of the gene, suggesting that the strength of the downstream effect associated with SF3B1 variants is proportional to their proximity to the hotspot (Figure S18E).

This approach also helped to explain why silent mutations were included in the best found subgrouping of NF1 in METABRIC(LumA). Although one should expect that these synonymous variants would have downstream effects equivalent to that of NF1 wild-types when compared to other types of NF1 point mutations, we found that a classifier trained to predict NF1 nonsense and silent mutations performed significantly better than the NF1 gene-wide classifier (21/48 NF1 point mutants, AUC=0.782 vs. AUC=0.660, down-sampled confidence=0.97) as well as the classifier trained to predict NF1 nonsense and missense mutations (31/48 NF1 point mutants, AUC=0.614). Because there were fewer than 20 samples total bearing NF1 nonsense mutations, no classifier was trained on them in isolation, which was also true of silent mutations.

Examining the distributions of the scores returned by these classifiers revealed that the combined NF1 nonsense and silent mutation subgrouping’s classifier was not only better able to distinguish between its own mutations and the remaining samples in the cohort, but it also did a better job of separating other NF1 mutations from NF1 wild-types than the gene-wide classifier, and especially NF1 missense mutations (Figure S19A). Furthermore, it successfully predicted the presence of silent mutations in held out samples. We thus conclude that these NF1 variants classified as synonymous very likely do have downstream transcriptomic effects aligned with those of active NF1 mutations. Although this finding contradicts the intuition that so-called silent mutations should not imbue significant downstream impacts, it is less surprising in light of prior work demonstrating that these mutations can indeed enact non-trivial effects on splicing, transcript folding/stability, translational rates, co-translational folding/stability, and degradation^53,54^. Although NF1 splice mutations are often mistaken for silent mutations by sequencing methods^55,56^, evidence that synonymous mutations of the NF1 gene are selected in cancers such as T-cell acute lymphoblastic leukemia^57^ signals a need for more research in this area.

An altogether different type of pattern was discovered within GATA3 in breast cancer where we found that GATA3 mutations not included in the best found subgroupings tended to have predicted scores between those assigned to samples carrying the subgrouping and GATA3 wild-types (Figures S19B-C). Further examination revealed that GATA3 mutations can be decomposed into disjoint pairs of subgroupings corresponding to whether they overlap with the zinc finger domain or the X308 splice site hotspot whose predicted scores were orthogonal to one another. This behavior was present in both METABRIC(LumA) and TCGA-BRCA(LumA), thus revealing the presence of two independent expression programs within GATA3 that are consistent across different breast cancer cohorts (Figure 5). This builds upon existing research demonstrating that GATA3 mutations can be partitioned into subsets with different functions and clinical outcomes by providing a transcriptomic characterization of these groupings^41,58^.

**FIG. 5.**
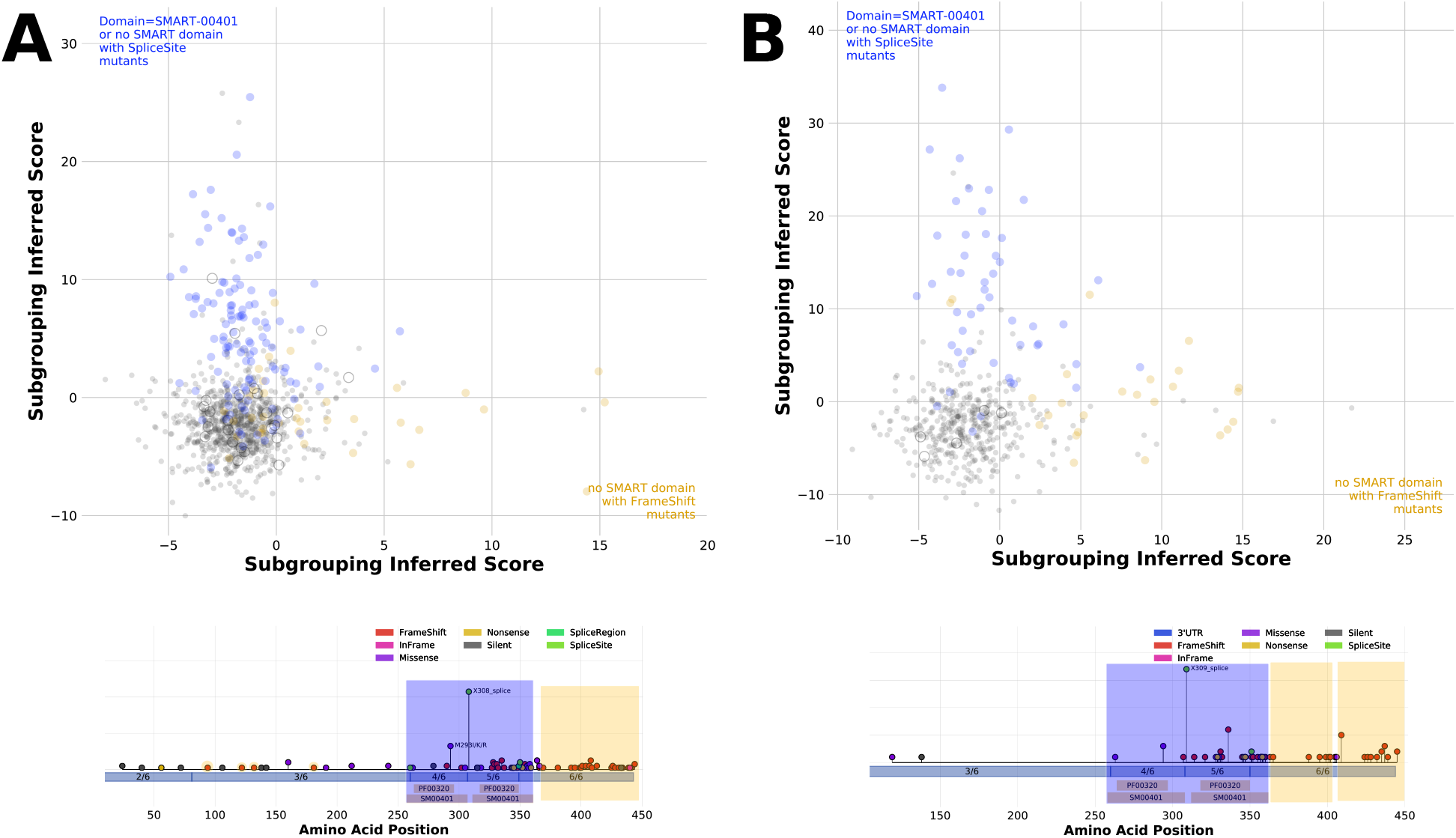
GATA3 downstream effects can be decomposed into two orthogonal axes. Amongst the divergent subgroupings enumerated for GATA3 in our breast cancer cohorts we found a pair of non-overlapping subgroupings that produced mutation scores with no correlation with one another in both METABRIC(LumA) (**A**) and TCGA-BRCA(LumA) (**B**). Each cohort sample is represented by a point, with samples shaded according to whether they carried a mutation in one of the subgroupings (blue/yellow fill), in neither (grey fill), or in both (hollow circle).

### Subgroupings enrich the characterization of drug response in cell lines

Do these divergences in downstream effects lead to divergent responses to pharmacological treatments? To answer that question, we tested the performance of our subgrouping classifiers in predicting response to drug interventions in cancer cell lines. We applied the classifiers we trained in each of our cohorts to the CCLE cohort^59^, which contains-omic and drug response data for 990 cell lines. For each classification task, we calculated the correlation between the mutation scores predicted for the CCLE cohort and drug response as measured by AUC50 for the 265 drugs which had response profiles available in at least 100 of the cell lines in the cohort where expression calls had also been made. We thus found that many subgroupings which exhibited divergent classification performance in the training cohort also yielded divergent associations with these clinical phenotypes.

For example, the scores returned by the best GATA3 subgrouping in METABRIC(LumA) consisting of mutations on the 5th exon and splice site mutations not on any exon consistently had stronger correlations with increased sensitivity to a wide range of drugs interrogated in CCLE, including tyrosine kinase inhibitors such as Dasatinib and Lapatinib (Figure 6A). The former is a selective SRC-family kinase inhibitor typically used for treatment of chronic myeloid leukemia and acute lymphoblastic leukemia^60^. Moreover, Dasatinib has also shown promise in breast cancer, with several clinical trials currently exploring its use as a monotherapy or in combination with other agents like Paclitaxel and Trastuzumab^61–63^. Zinc finger 2 (ZnFn2) mutations occur in the 5th exon of the GATA3 gene, and typically result in a truncated C-terminus. Because ZnFn2 mutations are known to cause observable decrease in typical GATA3 binding, these mutations are generally thought to cause loss of function^58^, however Znfn2 mutant GATA3 also has the potential to localize to a novel suite of target genes, resulting in increased expression at those sites^41^. ZnFn2 mutations are associated with very poor prognosis relative to other mutants, and cell lines harboring these mutants exhibit transcriptional reprogramming in favor of epithelial to mesenchymal transition (EMT)^41^. SRC and SRC-family kinases are also known to regulate EMT in solid tumors^64,65^, thus providing an indirect link to the observed sensitivity of cells harboring GATA3 Znfn2 mutations to Dasatinib.

**FIG. 6.**
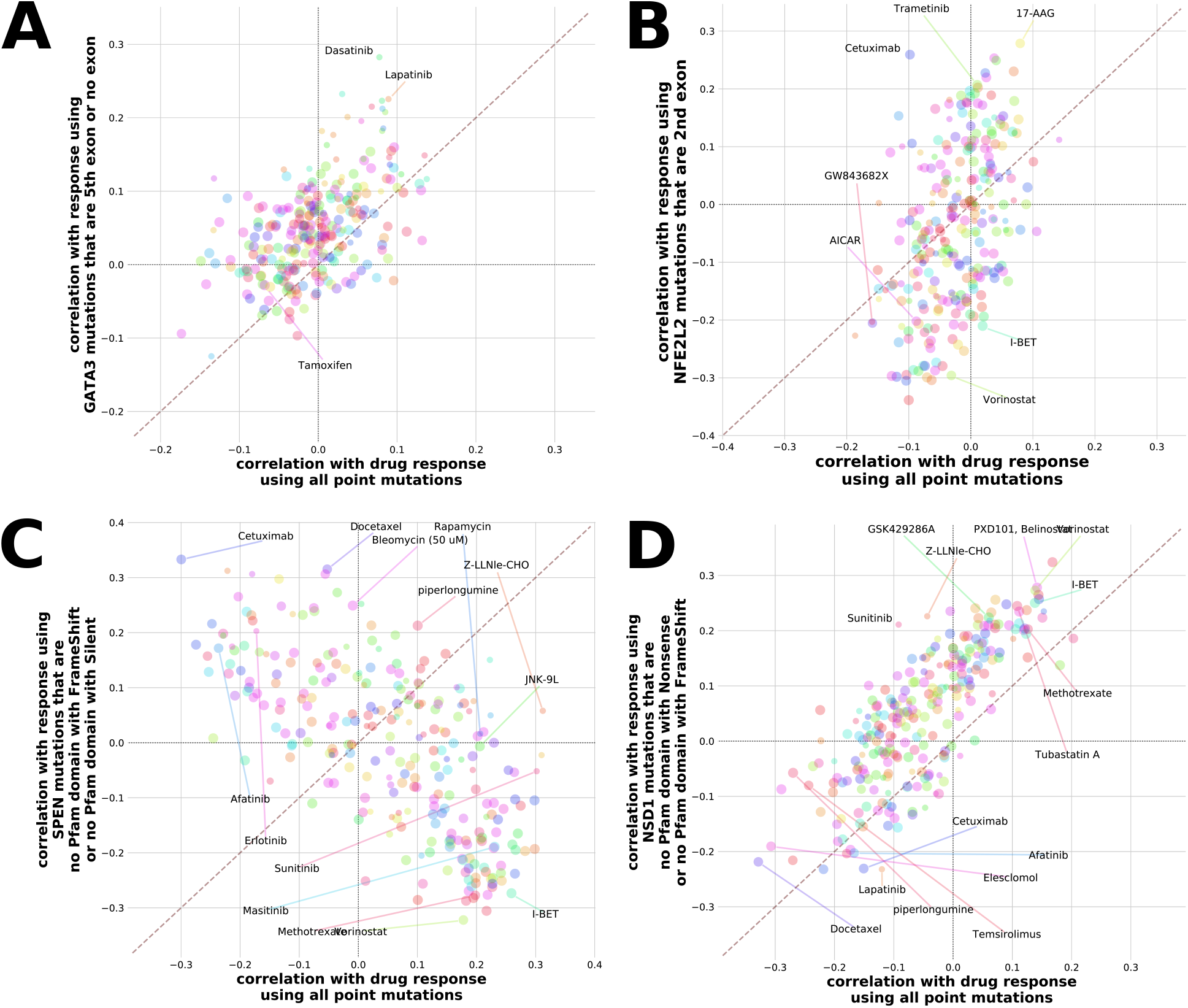
Using subgroupings improves concordance with clinically relevant phenotypes. We applied our trained classifiers to the CCLE cohort and computed the Spearman correlations between the scores returned by the classifiers and drug response for 265 compounds with AUC50s measured in at least 100 cell lines which also had expression calls available. For (**A**) GATA3 in METABRIC(LumA), NFE2L2 in TCGA-LUSC, (**C**) SPEN in TCGA-STAD, and (**D**) NSD1 in TCGA-HNSC(HPV-) we compared these correlations for the gene-wide classifier and the classifier of the best found subgrouping. Points correspond to individual drugs, with the area of each point proportional to the number of cell lines for which AUC50s were available for the given drug. Correlations were multiplied by −1, and thus higher correlations correspond to stronger association with increased sensitivity of the cell lines to the compound in question.

Other genes in which divergent subgroupings were associated with divergent drug response included NFE2L2 and SPEN (Figures 6B-C). In addition, we found cases such as NSD1 where the best found subgrouping was associated with a stronger overall association in cell lines across the majority of drugs included in this analysis (Figure 6D). These findings indicate that the subgroupings our approach discovers allow for the creation of transcriptomic models that are better able to characterize the impact of recurrent mutations on processes integral to tumorigenesis.

## DISCUSSION

We have introduced a method for exploring and characterizing the heterogeneity of alteration landscapes in genes frequently mutated in several cancers. In addition to ascertaining gene-wide transcriptomic signatures, this approach allowed us to systematically identify cancer genes containing subsets of mutations with functional effects that diverge from their remaining mutations. Considering subgroupings of mutations allowed us to find an expression profile associated with a gene implicated in tumorigenesis in many cases where it was otherwise difficult or impossible, and to discern which mutations in a gene are particularly likely to have a quantifiable downstream effect. The gains from considering a variety of mutation subgrouping tasks were far greater than from using more sophisticated classification algorithms, and often yielded more accurate models than using other methods to identify significant variants. Subgrouping signatures also exhibited strong performance in tumor cohorts to which they were not exposed during training, as well as improved association with drug response in cell lines.

Taken together, our findings confirm that genes with divergent mutation profiles are ubiquitous in cancer. Furthermore, they demonstrate that no characterization of the downstream effects of genes implicated in tumorigenesis is complete without taking these divergences into account. Our exploration of the subgrouping search space allowed us to construct more robust models linking the genome and transcriptome in tumor cohorts, as well as to predict the effects of mutations of unknown significance and to characterize the relationships between different perturbation axes extant within genes active in cancer processes. The detection of divergent alteration subgroupings has the potential to improve the specificity of precision treatments, aid in patient stratification, and to anticipate otherwise unexpected and undesirable therapeutic outcomes. Further, discovering subgroupings composed of mutations with convergent downstream effects may guide efforts to reposition existing pharmaceutical interventions to orthogonal scenarios that resemble approved clinical indications. This approach thus allows us to construct a more comprehensive catalogue of expression signatures associated with driver events in cancer, and illustrates that identifying subsets of mutations with unique transcriptomic signatures can yield robust and actionable biological insights.

## Supporting information

Graphical Abstract

## ACKNOWLEDGEMENTS

The authors would like to thank all members of the Pathways+Omics Group at OHSU for their support and suggestions related to this project, with particular gratitude to Joey Estabrook, Özgün Babur, Olga Nikolova, and Kevin Watanabe-Smith.

## SUPPLEMENTARY METHODS

### Cohort data preparation

We used a total of 18 publicly available tumor cohorts in this study, which included 15 individual cohorts from TCGA as well as METABRIC, Beat AML, and CCLE. These cohorts were selected on the basis of the availability of all three of expression, variant, and copy number data for the samples they contained (except for Beat AML, for which CNA calls were not made), as well as sufficient size (at least 200 samples with all three data types collected). Cohorts such as TCGA-COADREAD and TCGA-GBMLGG which are agglomerations of other cohorts were omitted.

For TCGA cohorts, Illumina RNAseq RSEM-normalized expression calls and GISTIC2.0 copy number calls were downloaded from the Broad Firehose portal (http://firebrowse.org/), while TCGA variant calls were downloaded from the Synapse portal for the MC3 pan-cancer analysis pipeline (https://www.synapse.org/#!Synapse:syn7214402). Expression, copy number, and variant data for the METABRIC and CCLE datasets were downloaded from cBioPortal (https://www.cbioportal.org/).

We applied UMAP (version 0.3.10) to project the expression profiles of the samples in each cohort into a two-dimensional space for easier interrogation of the global structures present within the transcriptome. These projections were compared against annotations of known molecular subtypes in cohorts where such annotations were available. We created sub-cohorts where UMAP transcriptome clusters overlapped with these subtypes (see Figures S1 and S2 and Table S1). Cases such as TCGA-SARC which initially passed the 200-sample threshold but had to be divided into sub-cohorts that did not meet the threshold were omitted from further analysis. Molecular subtype annotations for TCGA cohorts were provided by the Korkut Lab as part of the PCAWG Consortium; for METABRIC these annotations were downloaded from cBioPortal.

### Defining mutation subgroupings

Our mutation subgrouping method is based on organizing the genomic alterations present in a cohort according to various properties that mutations can have in common. The particular properties used in this study are:

- **Exon** The exon on which the mutation is located. The value ‘.’ was given to mutations such as splice site deletions which are not assigned to a specific exon. *e.g.* Exon = 5, Exon = 2
- **Amino Acid Location** The amino acid or acids affected by the mutation. The value ‘.’ was given to mutations for which this property is not applicable, such as intronic mutations. *e.g.* Location = 1047, Location = 274
- **Amino Acid Substitution** The specific protein substitution that takes place as a result of the mutation. *e.g.* H1047R, V600E
- **Form** The functional consequence of the mutation. *e.g.* Missense, Nonsense, FrameShiftIns, InFrameDel
- **Form(base)** The same as “Form”, but with insertions and deletions for a given type of mutation grouped together. For example, frameshift insertions and frameshift deletions are merged together into frameshifts. *e.g.* Missense, Nonsense, FrameShift, InFrame
- **SMART Domain** The SMART protein domain on which the mutation rests. Can also take on the value “no overlapping domain”. *e.g.* SM00233
- **Pfam Domain** The same as above but with Pfam protein domains. *e.g.* PF00853

All annotation levels except those related to protein domains were inferred from the MAF files listing the variant calls for each cohort. Protein domain data was downloaded from Ensembl (http://grch37.ensembl.org/downloads.html) using the following parameters, where *domain* refers to the SMART or Pfam databases as appropriate:

- `Dataset` Ensembl Genes 97 Human genes (GRCh37.p13)
- `Filters` Gene type: protein coding, With *domain* ID(s): Only
- `Attributes` Gene stable ID Transcript stable ID, *domain* ID, *domain* start, *domain* end

A subgrouping is thus defined by a nested combination of values chosen for one or more of these attributes. For example, a single hotspot mutation in PIK3CA can be represented by the subgrouping {*AAsub* = *H*1047*R*}. We can define the same subgrouping using additional properties: {*Exon* = 21 : *AAloc* = 1047 : *AAsub* = *H*1047*R*}. These additional properties are redundant in this case, as naturally all H1047R substitutions are located at amino acid 1047 and in turn all of the alterations at this amino acid are located on the 21st exon of PIK3CA. Nevertheless, we can expand this subgrouping to include other PIK3CA mutations which may or may not be functionally similar to H1047R. Thus we could consider the subgrouping {*Exon* = 21 : *AAloc* = 1047 : *AAsub* = (*H*1047*R or H*1047*L*)} to test the hypothesis that the particular amino acid that replaces the wildtype at this hotspot does not have an impact on downstream effects. Likewise, the subgroupings {*Exon* = 21 : (*AAloc* = 1047 : *AAsub* = *H*1047*R*) *or* (*AAloc* = 1049 : *AAsub* = *G*1049*R*)} and {*Exon* = 10 : *AAloc* = 542 : *AAsub* = *E*542*K or Exon* = 21 : *AAloc* = 1047 : *AAsub* = *H*1047*R*} can be used to compare hotspots at different loci within PIK3CA. We can also choose other properties to construct the same subgrouping based on which attributes of PIK3CA alterations we believe to be the most important in determining downstream effects: {*Form* = *Missense* : *AAsub* = *H*1047*R*}, {*Domain* = *SM* 00146 : *AAsub* = *H*1047*R*}, and so on.

### Enumeration of classification tasks in tumor cohorts

Cancer genes were identified using the OncoKB repository, with only genes included in at least one of the “Vogelstein”, “SANGER CGC(05/30/2017)”, “FOUNDATION ONE”, and “MSK-IMPACT” lists at https://oncokb.org/cancerGenes as of March 25th, 2019 being included for further analysis. In each cohort we considered the grouping of all point mutations in each such gene (referred to as the gene-wide task) and also sought to generate subgroupings of mutations within these genes.

We pruned the subgrouping search space by only using the four ordered mutation property hierarchies listed below, with the reasoning that a sizeable proportion of biologically relevant subgroupings of mutations could be generated using one of these combinations:

- `Exon` → `AA Location` → `AA Substitution`
- `Form` → `Exon`
- `SMART Domain` → `Form(base)`
- `Pfam Domain` → `Form(base)`

To further prune our search space, we only used subgroupings corresponding to a single branch containing at least twenty samples in one of these hierarchies as well as subgroupings corresponding to two branches each with at least ten samples. Branches did not have to terminate at a leaf node of the hierarchy. For example, using the combination `Form` → `Exon`, we could test {*PIK*3*CA* : *Missense*} as well as {*PIK*3*CA* : *Missense* : *Exon* = 10}, {*PIK*3*CA* : *Missense* : *Exon* = 21}, and {*PIK*3*CA* : *Missense* : *Exon* = 21 *or PIK*3*CA* : *Silent*}, but **not** {*PIK*3*CA* : *Missense* : *Exon* = (10 *or* 21) *or PIK*3*CA* : *Silent or PIK*3*CA* : *Nonsense*}.

To test the marginal benefit of relaxing this requirement, we also tested three-branch sub-groupings with at least five samples in each branch and twenty samples in total in the case of METABRIC(LumA). In all cases, subgroupings that contained all of the mutations of a gene in a cohort were discarded as being equivalent to the gene-wide task, which occurred in cases where the mutation hierarchy contained no more than two branches in total for a particular gene.

### Construction of classification tasks

A classification task was created for each of these enumerated subgroupings in a given cohort. To obtain a background distribution of predictive performance, we also added classification tasks using sets of samples randomly chosen from the cohort. Four such sets were created for each subgrouping already found, each of which contained the same number of samples as the number of samples carrying a mutation in the subgrouping in question. Two of these “random” subgroupings for each actual subgrouping chose samples from the entire cohort, while the other two only chose from the set of samples containing any point mutation in the gene mutated for the subgrouping.

Further classification tasks were added by considering copy number alterations as identified using discretized GISTIC 2.0 calls. For each of the non-random subgroupings described above, we created two new subgroupings by adding the set of samples carrying deep amplifications (+2) in the same gene as well as the set carrying deep deletions (−2). In cases where the given gene did not have at least five samples carrying the CNA to be added to the subgrouping, the corresponding subgrouping was excluded from further consideration. In genes where there were at least twenty deep amplifications or twenty deep deletions, we created a classification task containing just these CNAs of the gene.

Classification tasks were also constructed by dynamically discretizing PolyPhen and SIFT scores wherever these scores were available in the MAF file for the cohort (*i.e.* in TCGA cohorts). For each combination of mutated gene and variant significance metric, we enumerated all possible thresholds of the metric observed over variants of the gene in a cohort that yielded a unique subgrouping with at least twenty samples harbouring a mutation in the gene satisfying the threshold value (in the positive direction in the case of PolyPhen and the negative direction in the case of SIFT). For example, for AKT1 in TCGA-BRCA(LumA), we found the PolyPhen sub-groupings >= 0.006 and >= 0.999 (and no SIFT subgroupings), while for TP53 in TCGA-STAD we found 29 PolyPhen subgroupings (>= 0.002, >= 0.09, >= 0.275, …, >= 1.0) and 12 SIFT subgroupings (<= 0.8, <= 0.13, <= 0.11, …, <= 0).

### Training and evaluation of classifiers to identify transcriptomic signatures associated with subgroupings

Expression and variant data in each cohort was filtered to only include protein-coding genes on non-sex chromosomes prior to classifier training. Remaining expression data was then filtered to exclude gene features in the bottom five percentiles according to average value across the cohort before being log-normalized and then scaled using z-scores for each genetic feature. In each task we further excluded expression features associated with genes on the same chromosome as the gene containing the task’s subgrouping.

Each classification task consisted of predicting a vector of binary mutation labels using this processed expression matrix for a given cohort. The label for each sample in a task was ‘True’ if and only if it harboured any mutation within the subgrouping, or if it was randomly chosen from the set of cohort samples or the set of gene mutants as applicable for random background subgroupings. Predictions were made using the following algorithms implemented in scikit-learn (version 0.21.2), with any parameters not explicitly listed above being set to the default value:

- **Ridge Regression** `sklearn.linear model.LogisticRegression`

– tuning over eight values of *C*: [10^−7^, 10^6^, 10^5^, …, 10^0^]
– solver = ‘liblinear’, penalty = ‘l2’, max iter = 200, class weight = ‘balanced’
- **Support Vector Machine** `sklearn.svm.SVC`

– tuning over eight values of *C*: [10^−3^, 10^−2^, 10^−1^, …, 10^4^]
– kernel = ‘rbf’, gamma = ‘scale’, probability = True, cache size = 500, class weight = ‘balanced’
- **Random Forests** `sklearn.ensemble.RandomForestClassifier`

– tuning over eight values of *min samples leaf* : [1, 2, 3, 4, 6, 8, 10, 15]
– n estimators = 5000, class weight = ‘balanced’
- **Ridge Regression (deeper tuning)** `sklearn.linear model.LogisticRegression`

– tuning over *C* = [10^−8.2^, 10^−7.8^, 10^7.4^, …, 10^4.2^]
– solver = ‘liblinear’, penalty = ‘l2’, max iter = 200, class weight = ‘balanced’

Forty classifiers were fit for each task, corresponding to ten iterations of 4-fold cross-validation. The samples in each cohort were partitioned into quarters at random ten times; each classifier was thus tuned and trained on three such quarters before being asked to make predictions on the remaining quarter of samples. The same forty training and testing sub-cohorts were used across all tasks on a given cohort.

Each iteration of a classifier was tuned by training the classifier using each of the values in the classifier’s hyper-parameter tuning grid on four randomly-chosen subsets consisting of 80% of the training sub-cohort. The accuracy for each hyper-parameter tuning value was measured by taking the worst AUC across its four trained classifiers on the remaining 20% of the samples in the training sub-cohort. The best hyper-parameter value according to this metric was then used when training the classifier on the entire training sub-cohort before applying it to the entire testing sub-cohort. Classifier task performance on these testing sub-cohorts was measured using AUC as calculated by averaging predicted mutation scores for each cohort sample from all ten folds, segregating scores for mutated samples and wild-type samples, and then calculating the probability that a randomly-chosen mean score for a mutated sample was greater than a randomly chosen mean score for a wild-type sample across all possible such sample pairs.

We also created a bootstrapped population of downsampled AUCs by building a subset of samples to which each sample in the cohort was assigned to with probability *p* = 0.5, recalculating the AUC using the same method as above using just this subset, and repeating the process 500 times. This allowed us to estimate the robustness of our original AUCs with respect to variance in the training population. It also allowed us to calculate a score measuring the confidence that an AUC measured for one task was higher than that of another by calculating the probability that a downsampled AUC for one task was higher than a downsampled AUC for the other task over all 250,000 possible such pairs. Thus two equivalent tasks would be expected to have a confidence score of 0.5 in either direction, while a task with a significantly higher/lower AUC than another would be expected to have a confidence score of 1/0 compared to the other. This “AUC of AUCs” was especially useful when comparing the optimal subgrouping task found for a gene to its gene-wide counterpart. The same 500 subsets of samples were used to construct this confidence score in all classification tasks in a given training cohort.

Classifier task performance was further measured on each of the cohorts other than the one the classifier was trained on by applying each of the forty trained classifier iterations to their processed expression data. AUCs for these “transfer” experiments were calculated using the same sample-average method as described in the within-cohort case, this time with forty classifier output values for each sample.

### Measuring concordance between subgrouping classifier output and drug response

Summaries of cell line drug response observed within the CCLE cohort as measured by AUC50 were extracted from Table S4B downloaded from https://www.cancerrxgene.org/gdsc1000/GDSC1000_WebResources/Home.html. Subgrouping classifiers trained on TCGA and METABRIC cohorts were asked to make predictions for the CCLE cohort in the same manner as described for the transfer experiment above. For each combination of drug and task, we thus measured a correlation between subgrouping classifier output and drug response by calculating the Spearman rho between the AUC50 values and the average classifier predictions across the subset of samples for which drug response was available.

### Data and code availability

All of the code used for data preparation, machine learning analysis, and plot creation was written in Python 3.6 and can be found at https://github.com/ohsu-comp-bio/dryad and https://github.com/ohsu-comp-bio/HetMan. Assistance with the reproduction of the results presented herein will be provided upon request.

## SUPPLEMENTARY FIGURES AND TABLES

**FIG. S1.**
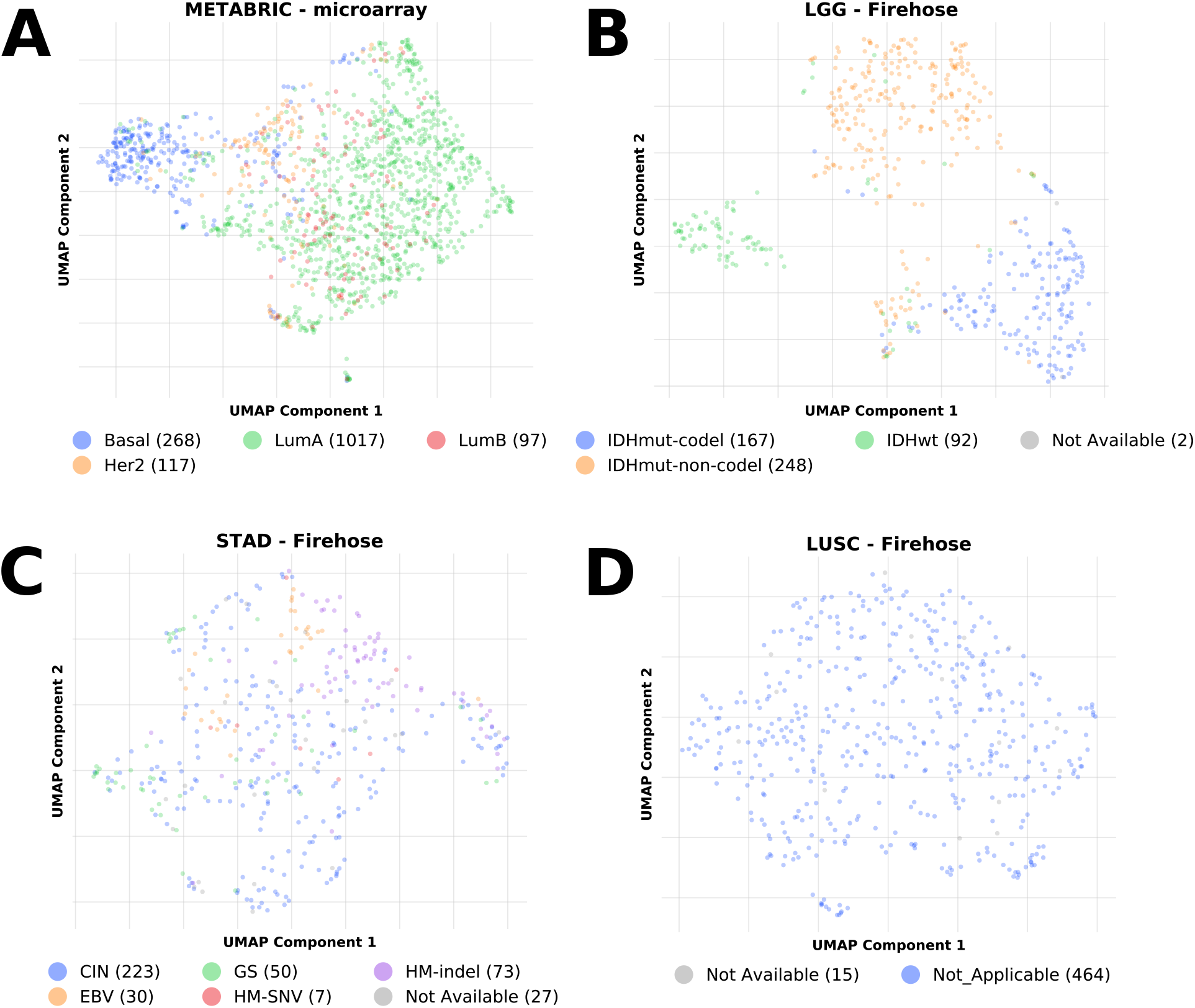
Clustering of cohort transcriptomes reveals profiles consistent with molecular subtypes. We applied unsupervised learning to the expression data used for each cohort considered in this study in order to remove unwanted variation associated with molecular subtypes from our alteration divergence analysis. In conjunction with information on known molecular subtypes present in these cohorts, we identified cases such as (**A**) METABRIC and (**B**) TCGA-LGG in which these subtypes clearly overlapped with distinguishable transcriptomic profiles. This contrasted with cohorts such as (**C**) TCGA-STAD in which subtypes were present but could not be unambiguously linked with unique transcriptomic profiles, and those like (**D**) TCGA-LUSC in which neither molecular subtypes nor expression clusters were present. The counts of cohort samples with each subtype are listed in the subplot legends.

**FIG. S2.**
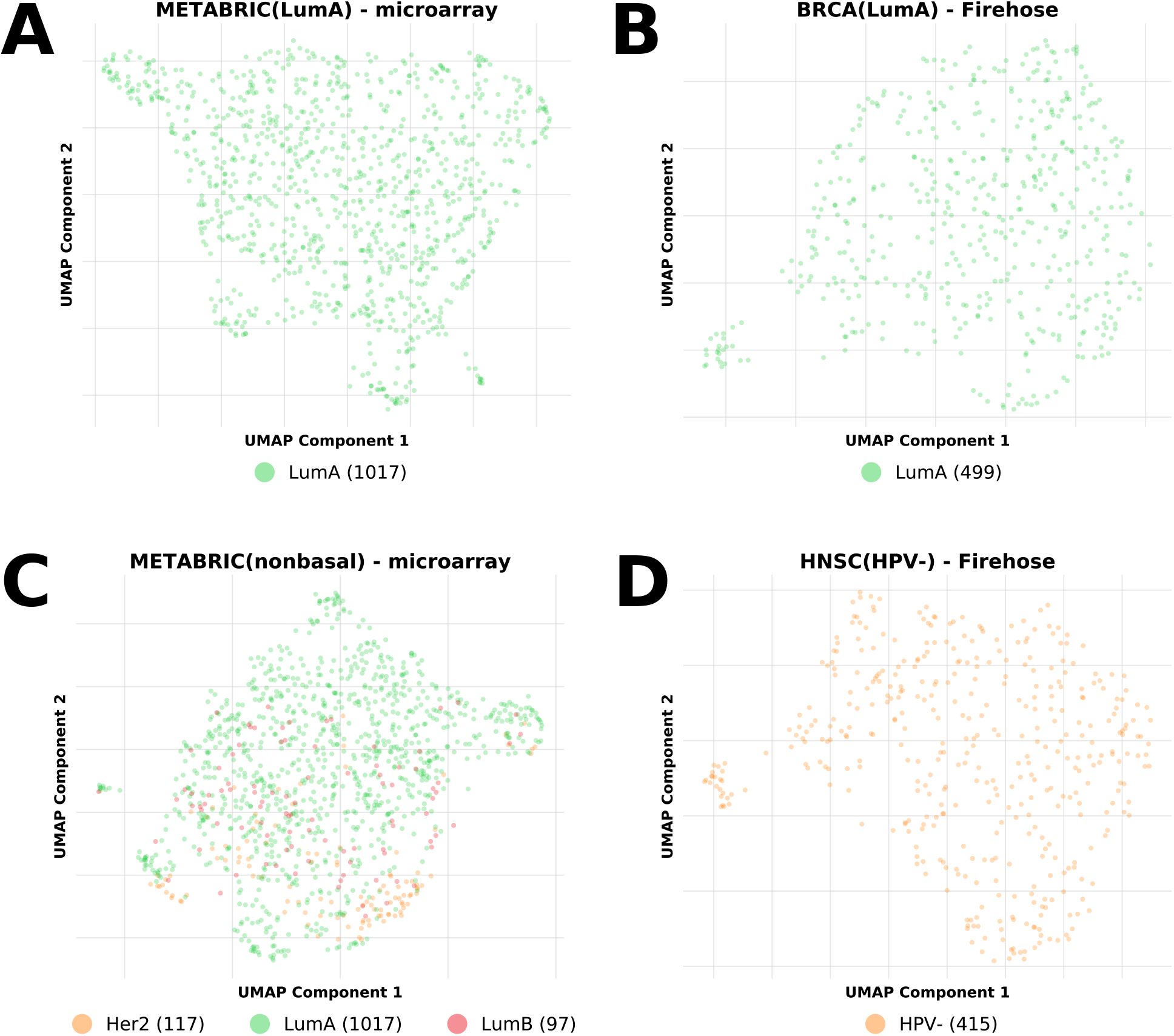
Subdividing cohorts yields more uniform training expression data. In cohorts where molecular subtypes were found to have identifiable transcriptomic profiles we created sub-cohorts that only included samples from a particular subtype or set of subtypes. Unsupervised learning on these sub-cohorts’ transcriptomes revealed that they did not exhibit the large-scale clusters of samples observed in the original cohorts and were thus much more suitable as input for our mutation classification pipelines.

**TABLE S1.**
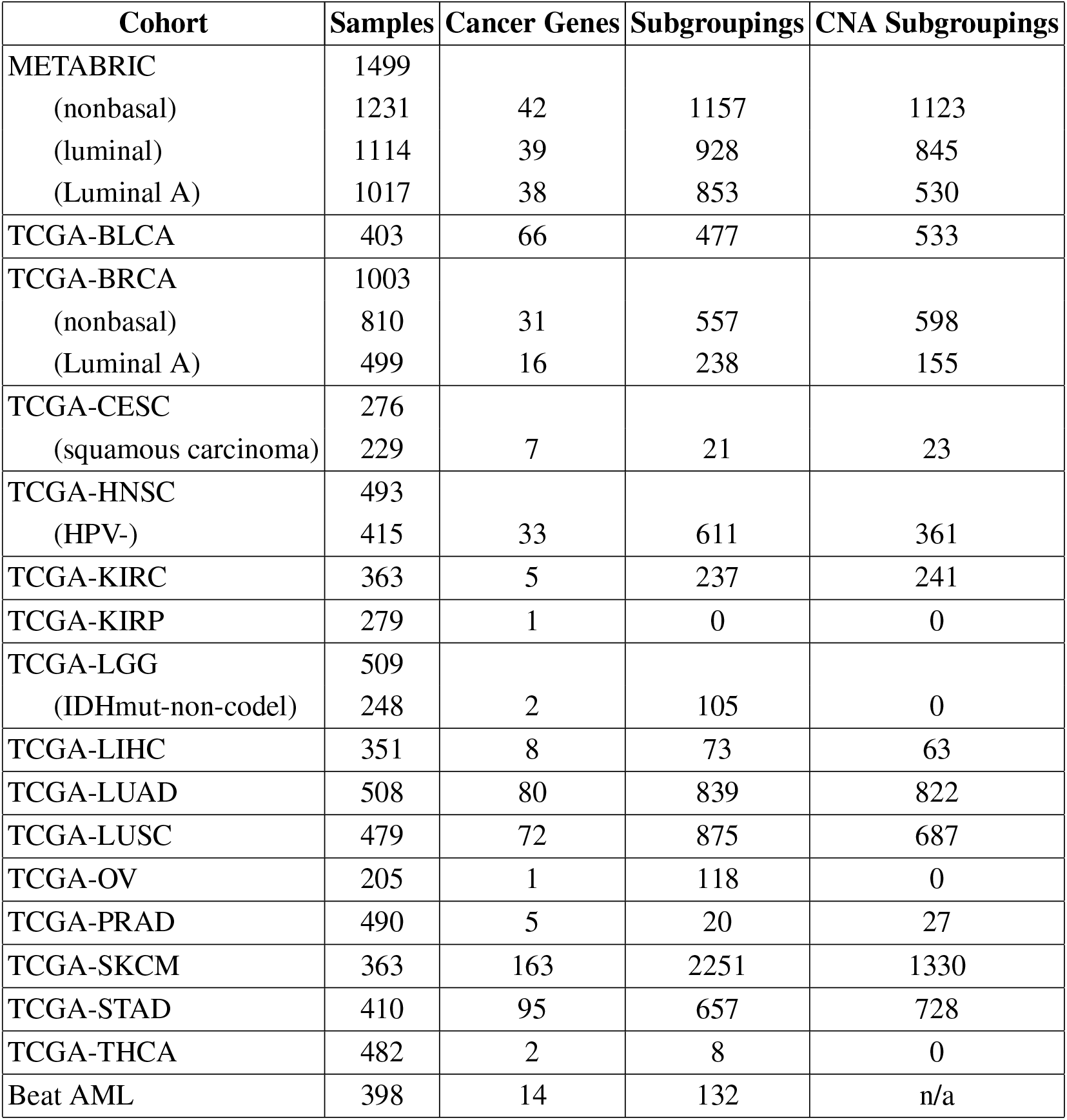
Cohorts used for mutation classifier training. In each cohort meeting our selection criteria we found the cancer genes with point mutations in at least twenty samples and enumerated their mutation subgroupings. Sub-cohorts identified using molecular subtypes and unsupervised learning are listed where applicable.

| **Cohort** | **Samples** | **Cancer Genes** | **Subgroupings** | **CNA Subgroupings** |
| --- | --- | --- | --- | --- |
| METABRIC | 1499 |  |  |  |
| (nonbasal) | 1231 | 42 | 1157 | 1123 |
| (luminal) | 1114 | 39 | 928 | 845 |
| (Luminal A) | 1017 | 38 | 853 | 530 |
| TCGA-BLCA | 403 | 66 | 477 | 533 |
| TCGA-BRCA | 1003 |  |  |  |
| (nonbasal) | 810 | 31 | 557 | 598 |
| (Luminal A) | 499 | 16 | 238 | 155 |
| TCGA-CESC | 276 |  |  |  |
| (squamous carcinoma) | 229 | 7 | 21 | 23 |
| TCGA-HNSC | 493 |  |  |  |
| (HPV-) | 415 | 33 | 611 | 361 |
| TCGA-KIRC | 363 | 5 | 237 | 241 |
| TCGA-KIRP | 279 | 1 | 0 | 0 |
| TCGA-LGG | 509 |  |  |  |
| (IDHmut-non-codel) | 248 | 2 | 105 | 0 |
| TCGA-LIHC | 351 | 8 | 73 | 63 |
| TCGA-LUAD | 508 | 80 | 839 | 822 |
| TCGA-LUSC | 479 | 72 | 875 | 687 |
| TCGA-OV | 205 | 1 | 118 | 0 |
| TCGA-PRAD | 490 | 5 | 20 | 27 |
| TCGA-SKCM | 363 | 163 | 2251 | 1330 |
| TCGA-STAD | 410 | 95 | 657 | 728 |
| TCGA-THCA | 482 | 2 | 8 | 0 |
| Beat AML | 398 | 14 | 132 | n/a |

**FIG. S3.**
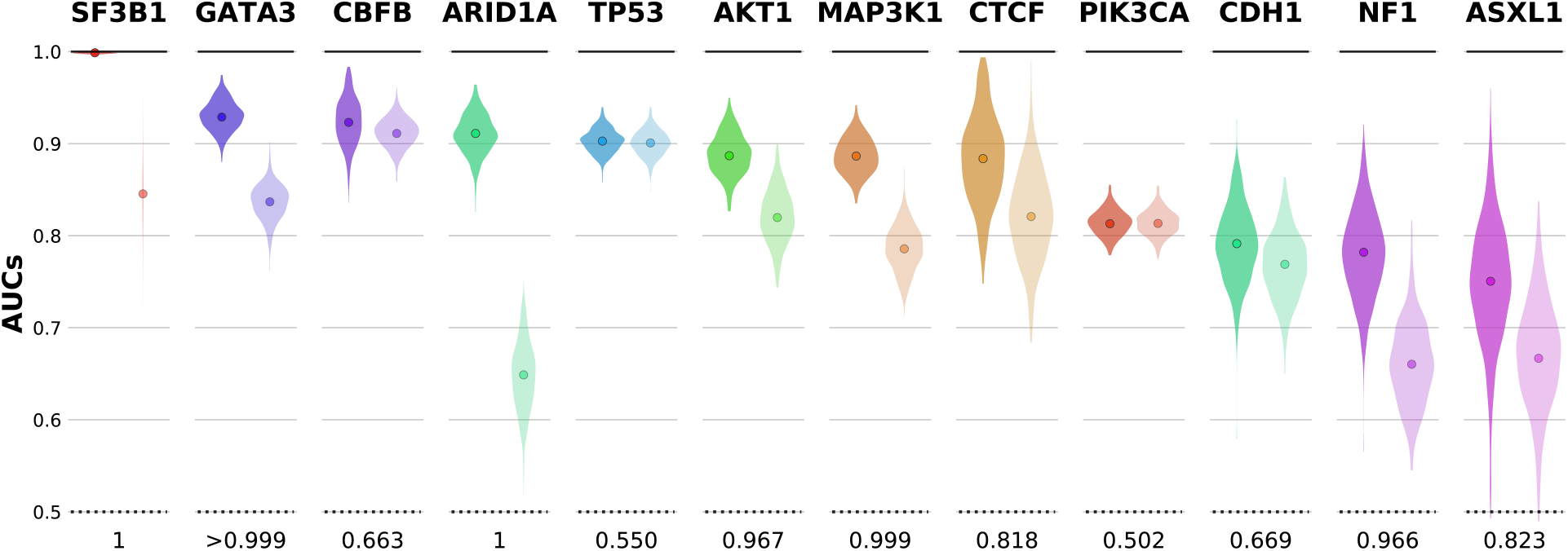
Down-sampling derives confidence intervals for classification task AUCs. For each prediction task we created a pool of 500 AUCs recalculated using randomly-chosen subsets of samples in the given cohort. These populations of down-sampled AUCs allowed us to estimate the likelihood that the classification task associated with a subgrouping yielded better performance than the task associated with all point mutations of its parent gene. For example, the down-sampled AUCs for the best found subgrouping within genes identified as exhibiting divergence in METABRIC(LumA) (darker violin-plots) tended to be higher than those recorded for the gene-wide tasks (lighter violins). The points within each violin denote the original AUC measured for the task using all samples in the cohort. Each gene’s panel is annotated with the probability that a down-sampled AUC for its best found subgrouping is greater than a down-sampled AUC for its gene-wide task.

**FIG. S4.**
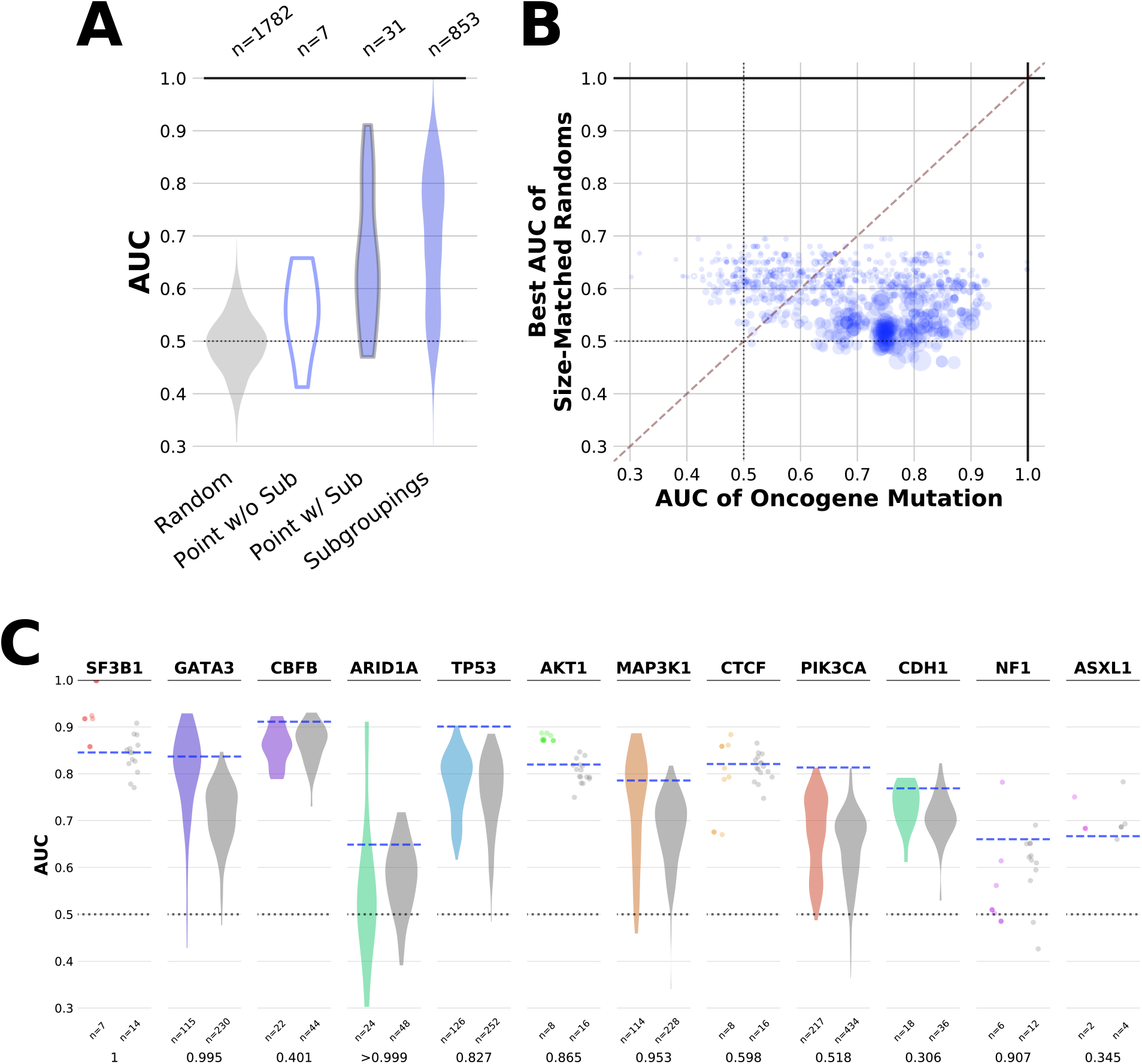
Subgrouping prediction tasks outperform cohort-specific and gene-specific random background prediction tasks. For each of the learning tasks completed in METABRIC(LumA) to predict the presence of an actual mutation or subgrouping of mutations, four additional tasks were performed based on predicting a simulated set of point mutations. (**A**) Two of these sets were constructed by randomly selecting a group of samples from the entire cohort of the same size as the group of samples affected by the original “real” mutation in the cohort. Tasks associated with actual mutations had higher AUCs as a population than tasks associated with cohort-specific random sets, and also when compared to random tasks with the same number of samples in the mutated set (**B**). (**C**) The other two sets were constructed by randomly selecting size-matched groups from the set of samples carrying any point mutation of the gene in the cohort. The AUCs of these gene-specific tasks (grey distributions) tended to be lower than the AUCs of subgroupings in genes that had been found to exhibit alteration divergence (colorful distributions). The probability that a down-sampled AUC for the best found subgrouping was greater than a down-sampled AUC for the best-performing of these random tasks created for each gene is listed below the pair of distributions plotted for each gene.

**FIG. S5.**
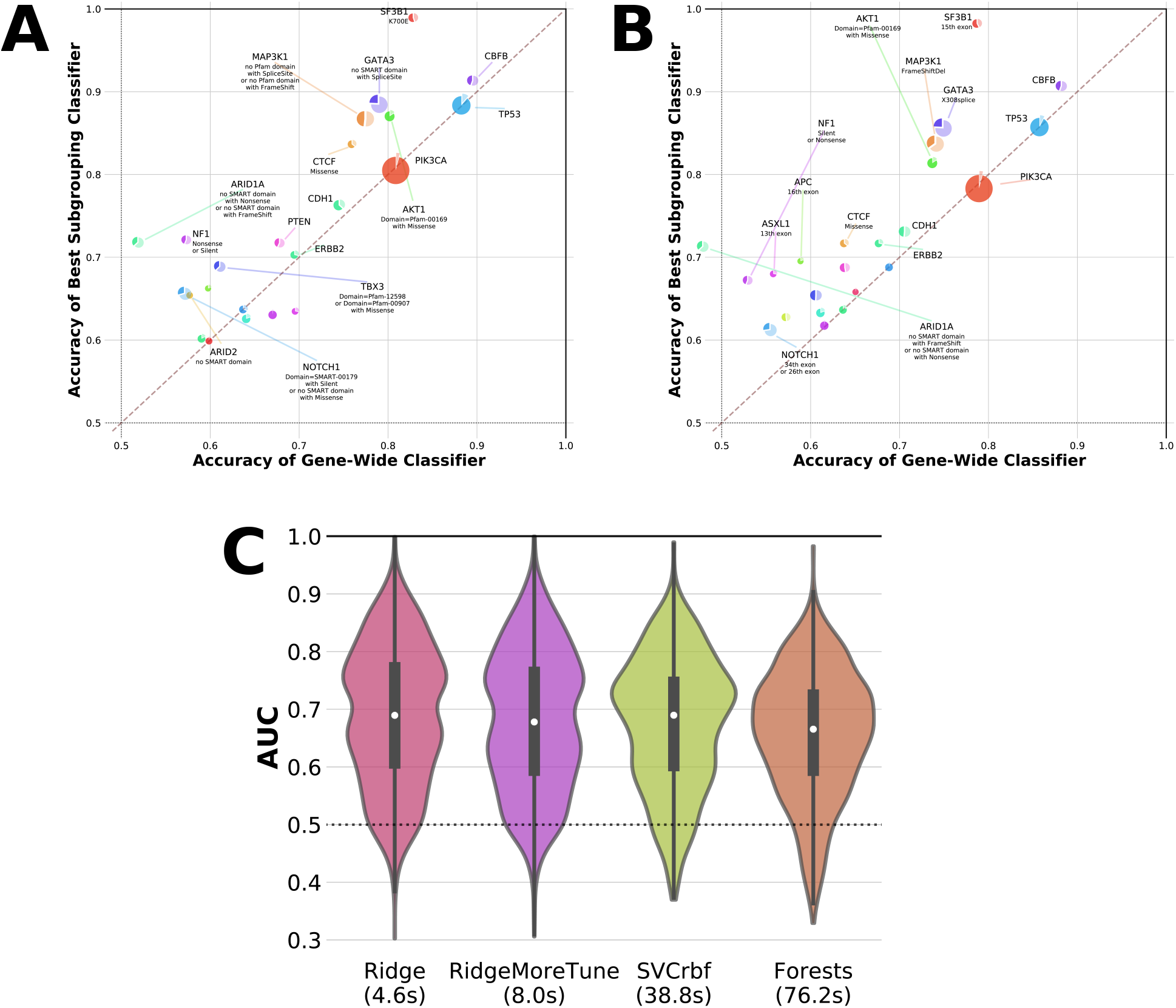
Increasing computational complexity does not change or improve upon classification performance. We observed similar subgrouping classification performance in METABRIC(LumA) when we repeated our prediction tasks with (**A**) a support vector machine classifier and (**B**) a random forest classifier in place of the ridge regression classifier that was originally used. (**C**) Using these more computationally complex classifiers did not result in improved classification performance across all non-random classification tasks.

**FIG. S6.**
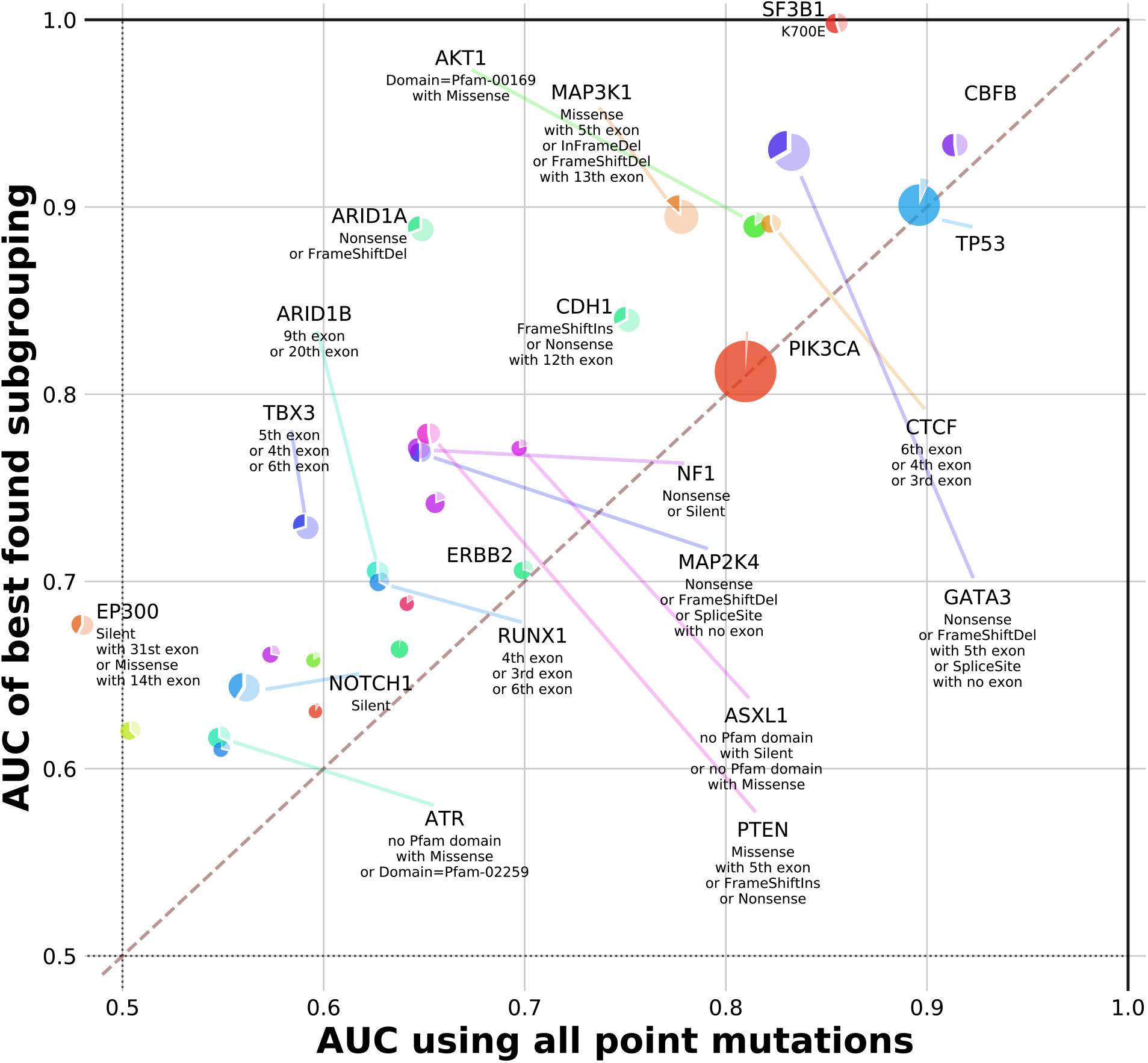
Applying an expanded search space to subgrouping enumeration and classification in METABRIC(LumA). The task enumeration step in METABRIC(LumA) was modified to allow for subgroupings of up to three branches each containing at least five samples for a total of at least twenty samples. This resulted in an expanded search space of 7598 subgroupings. The AUCs of the optimal subgroupings found for each gene are shown here in the same style as in Figure 1.

**FIG. S7.**
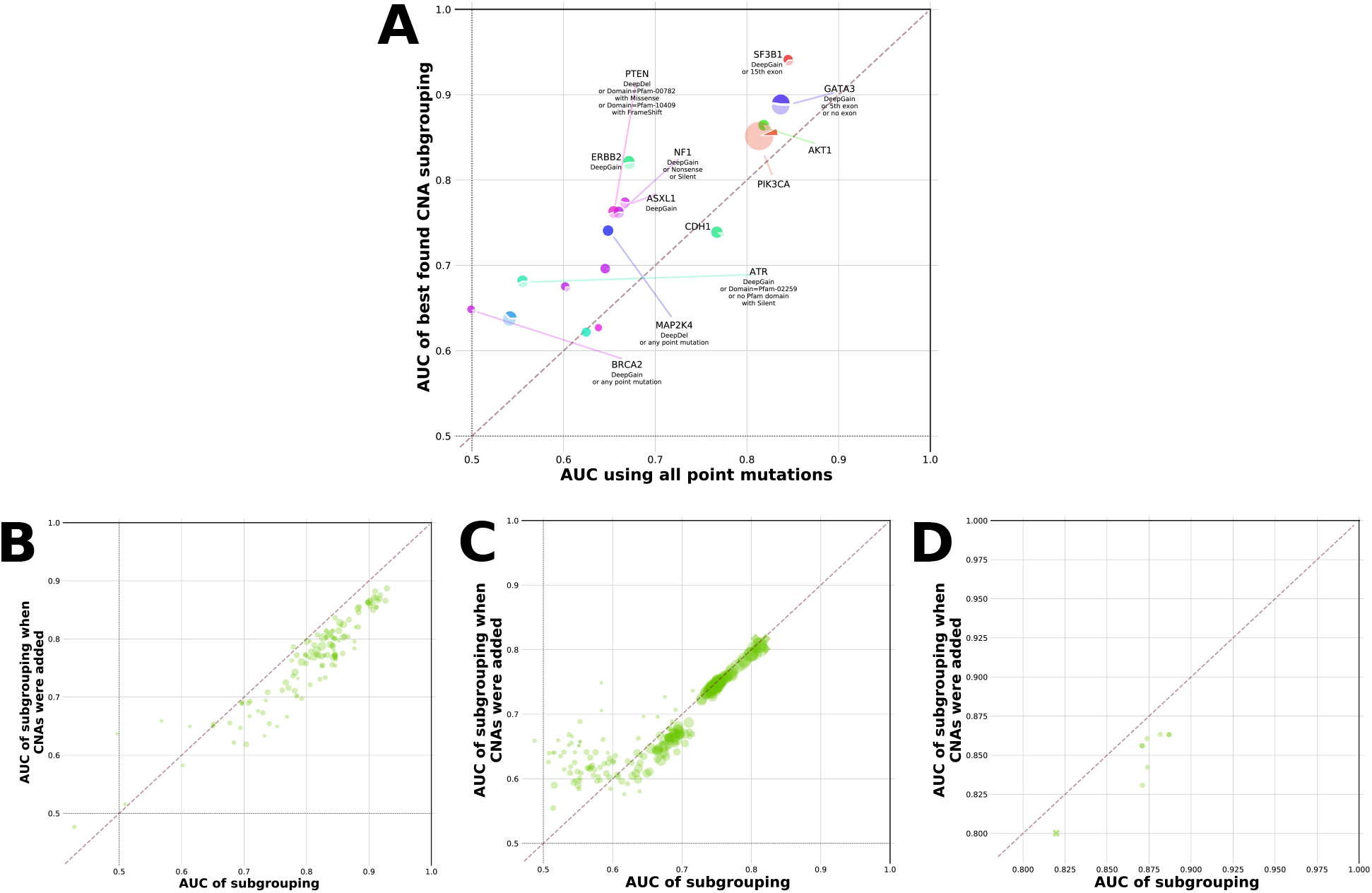
Adding copy number alterations to subgrouping classifiers in METABRIC(LumA). We augmented our classification tasks in METABRIC(LumA) by adding deep amplifications and deep deletions to each subgrouping where there were at least five of one of these two types of mutations present in the corresponding gene within the cohort. (**A**) Comparing the classification performance of the best found subgrouping containing CNAs (y-axis) to the gene-wide task (x-axis) for each cancer gene with enumerated subgroupings in METABRIC(LumA). (**B-D**) Comparing the performance for each subgrouping originally enumerated for GATA3, PIK3CA, and AKT1 respectively in METABRIC(LumA) (x-axes) to the performance of the same subgrouping combined with deep amplifications (y-axes). The point corresponding to the gene-wide classifier is marked with an ‘X’ in each panel.

**FIG. S8.**
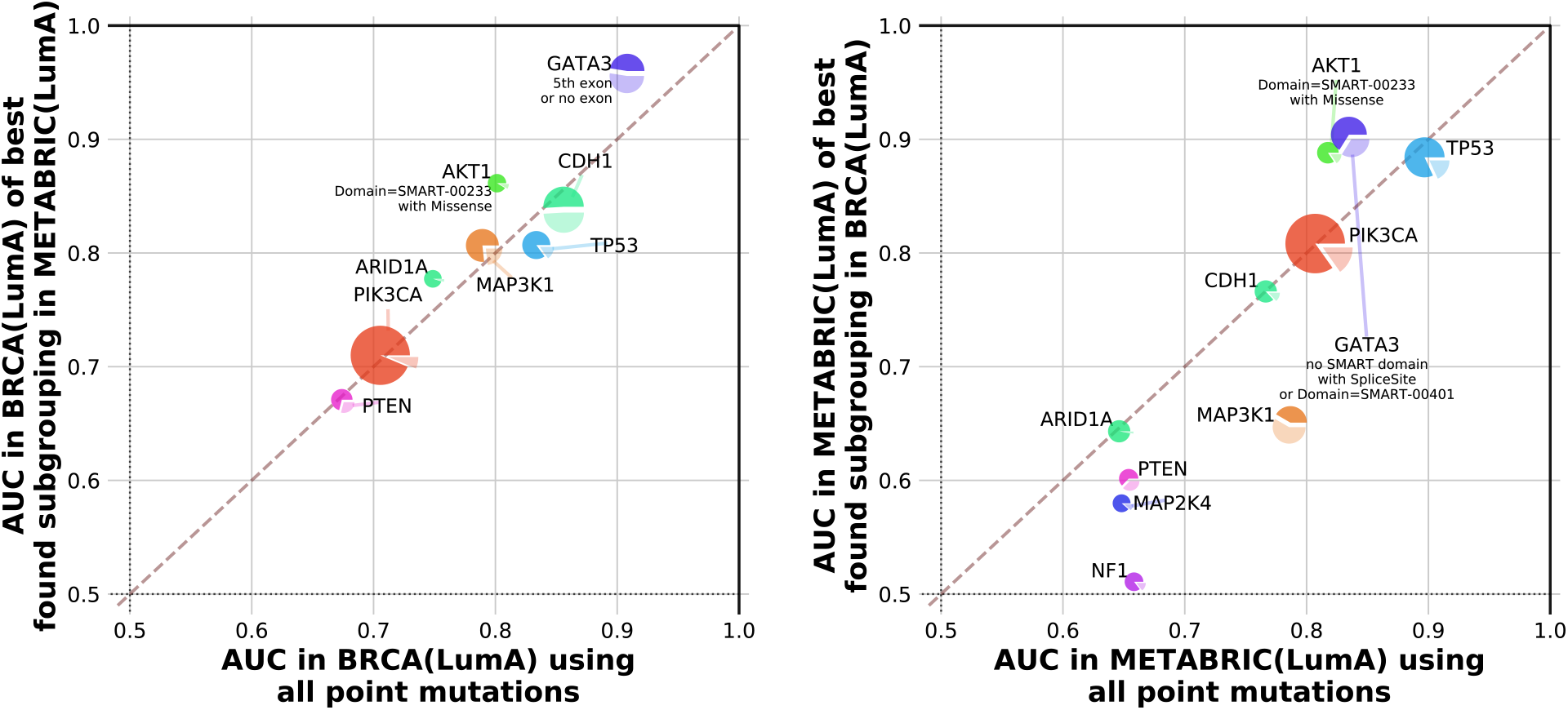
Relative performance of subgrouping tasks replicates across breast cancer cohorts. We compared the best-performing subgroupings found in METABRIC(LumA) to their corresponding gene-wide tasks using AUCs measured when training and testing classifiers in TCGA-BRCA(LumA) (left) and vice versa (right). This revealed that the particular subgroupings found to be most divergent tended to be consistent between different cohorts from the same cancer context.

**FIG. S9.**
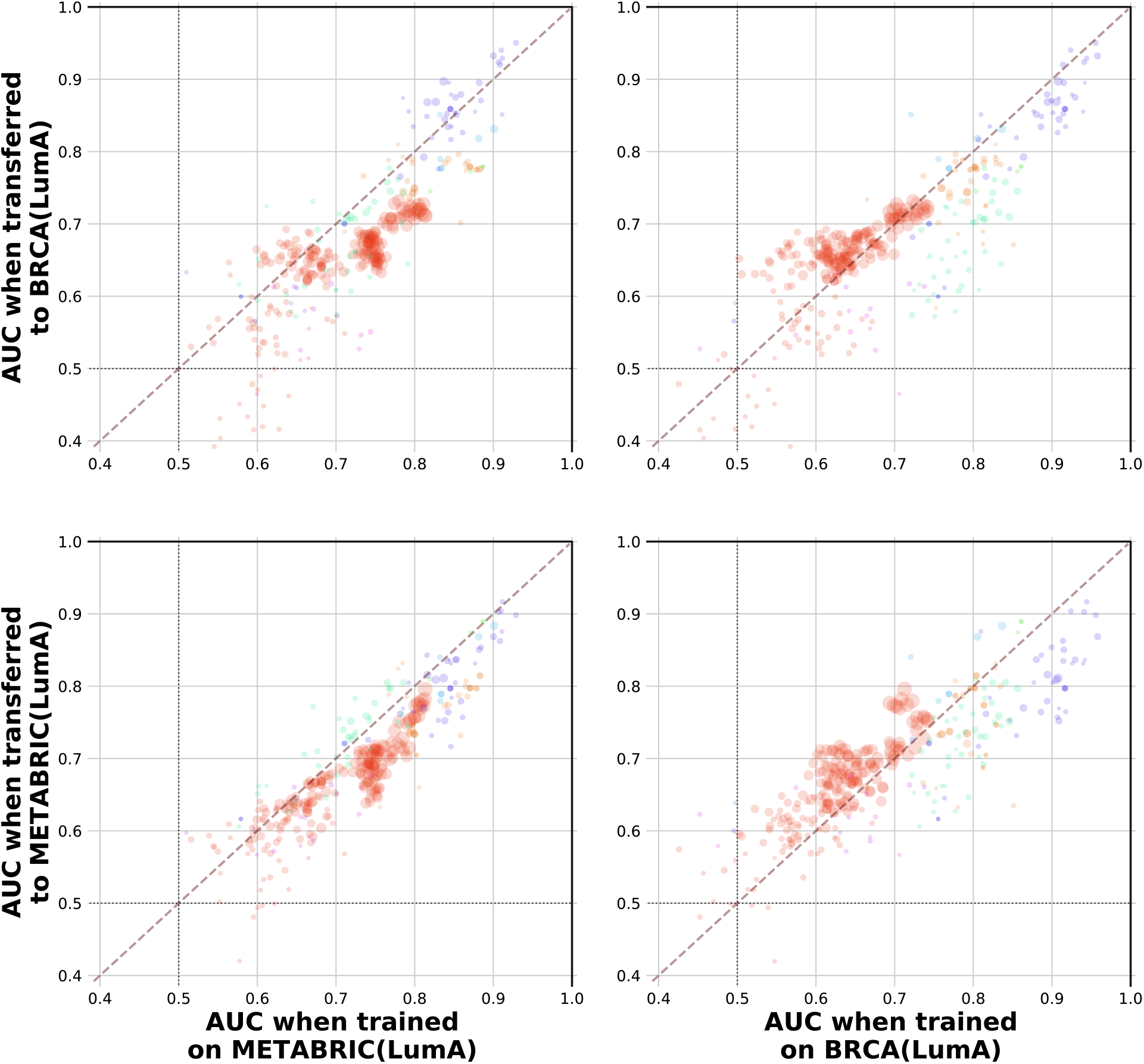
Subgrouping classification tasks preserve their efficacy when transferred across breast cancer cohorts. We asked the linear regression models trained to predict mutation subgroupings in METABRIC(LumA) to make predictions using the TCGA-BRCA(LumA) expression data (top row), and likewise using trained TCGA-BRCA(LumA) models and METABRIC(LumA) expression data (bottom row). These transferred models were successful in recapitulating their performance relative to that which was observed in the cohort within which they were trained (top-left and bottom-right) and relative to that which observed by models trained on the cohort they were transferred to (bottom-left and top-right). Points in each panel correspond to individual classification tasks, with colors chosen according to the mutated gene using the same color scheme as above, and point areas proportional to the geometric mean of the frequency of the mutation across the two cohorts as in Figure 2A.

**FIG. S10.**
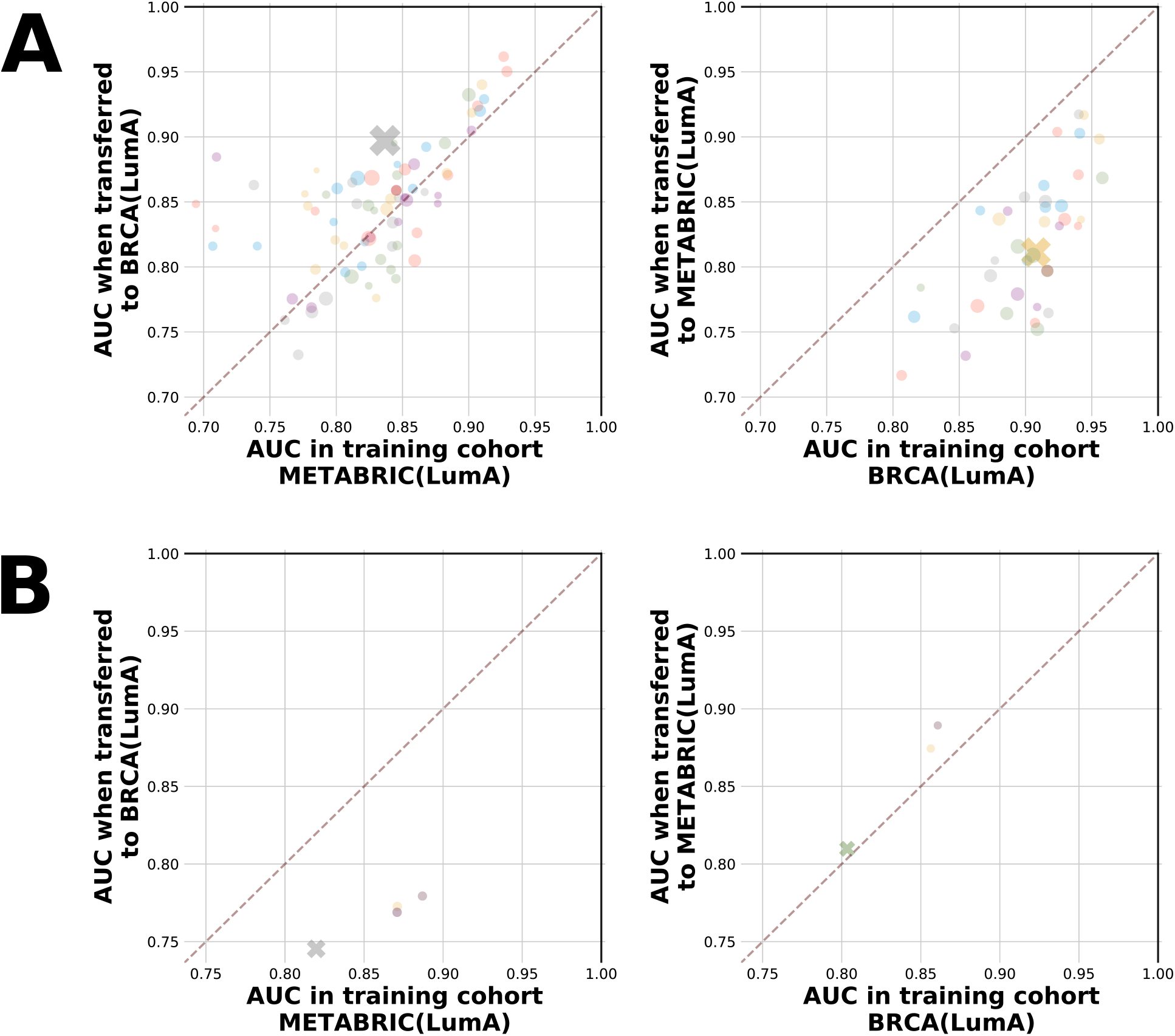
Subgrouping divergence is preserved when classifiers are transferred across breast cancer cohorts. The performance of mutation classifiers trained to predict (**A**) GATA3 mutations and (**B**) AKT1 mutations in the METABRIC(LumA) and TCGA-BRCA(LumA) cohorts was measured both in the original training cohort and when transferred to the other breast cancer cohort. Points correspond to individual classification tasks, with the gene-wide task highlighted with a colored ‘X’ in each panel, and point areas proportional to the frequency of the mutation in the training cohort.

**FIG. S11.**
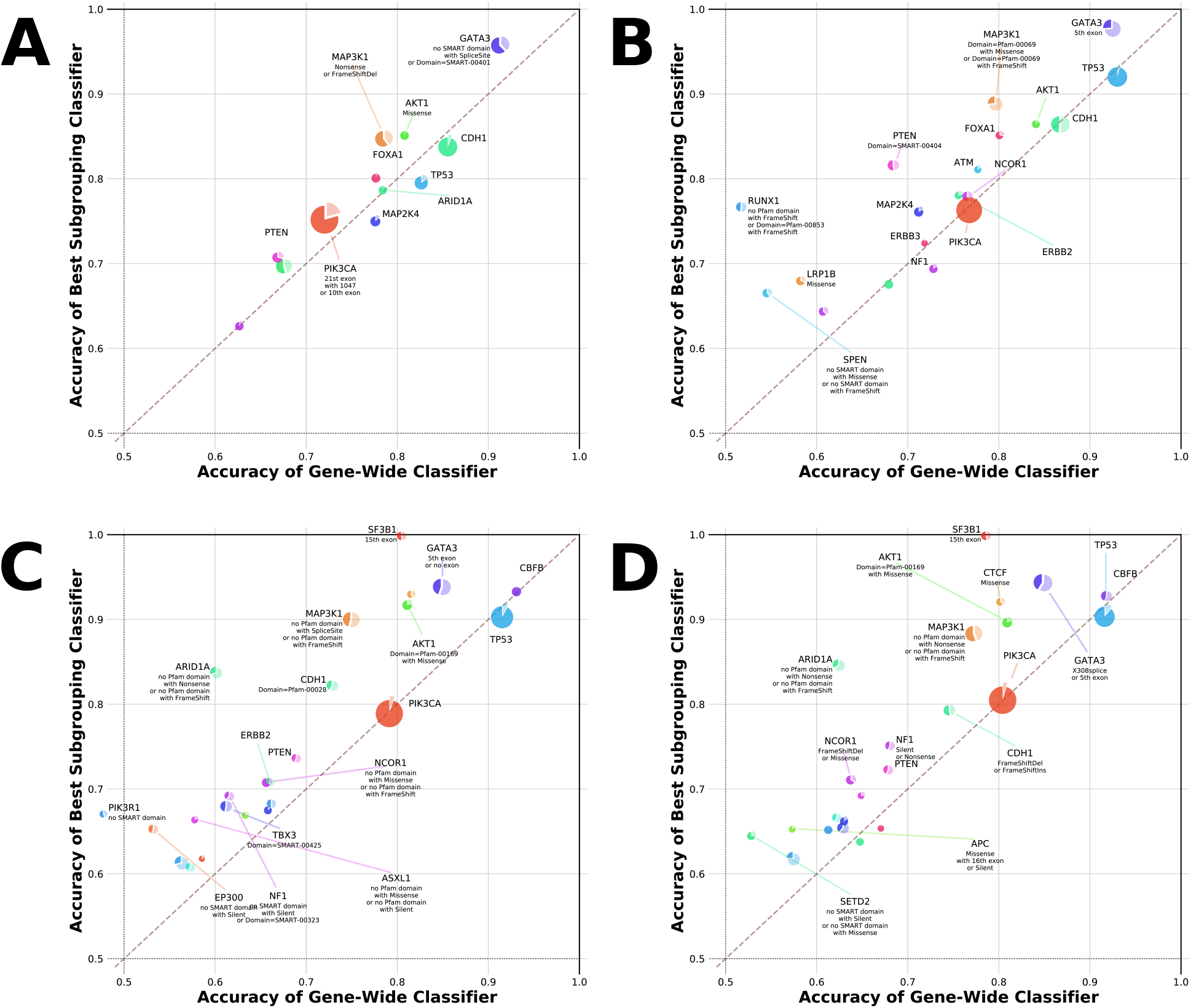
Subgrouping behaviour replicates across various choices of breast cancer expression datasets. We observed subgrouping classification performance similar to that in METABRIC(LumA) and TCGA-BRCA(LumA) when we repeated our prediction tasks using (**A**) kallisto TPM expression calls instead of Firehose RSEMs in TCGA-BRCA(LumA), (**B**) all nonbasal subtypes present in TCGA-BRCA, (**C**) all nonbasal subtypes present in METABRIC, and (**D**) both luminal subtypes present in METABRIC.

**FIG. S12.**
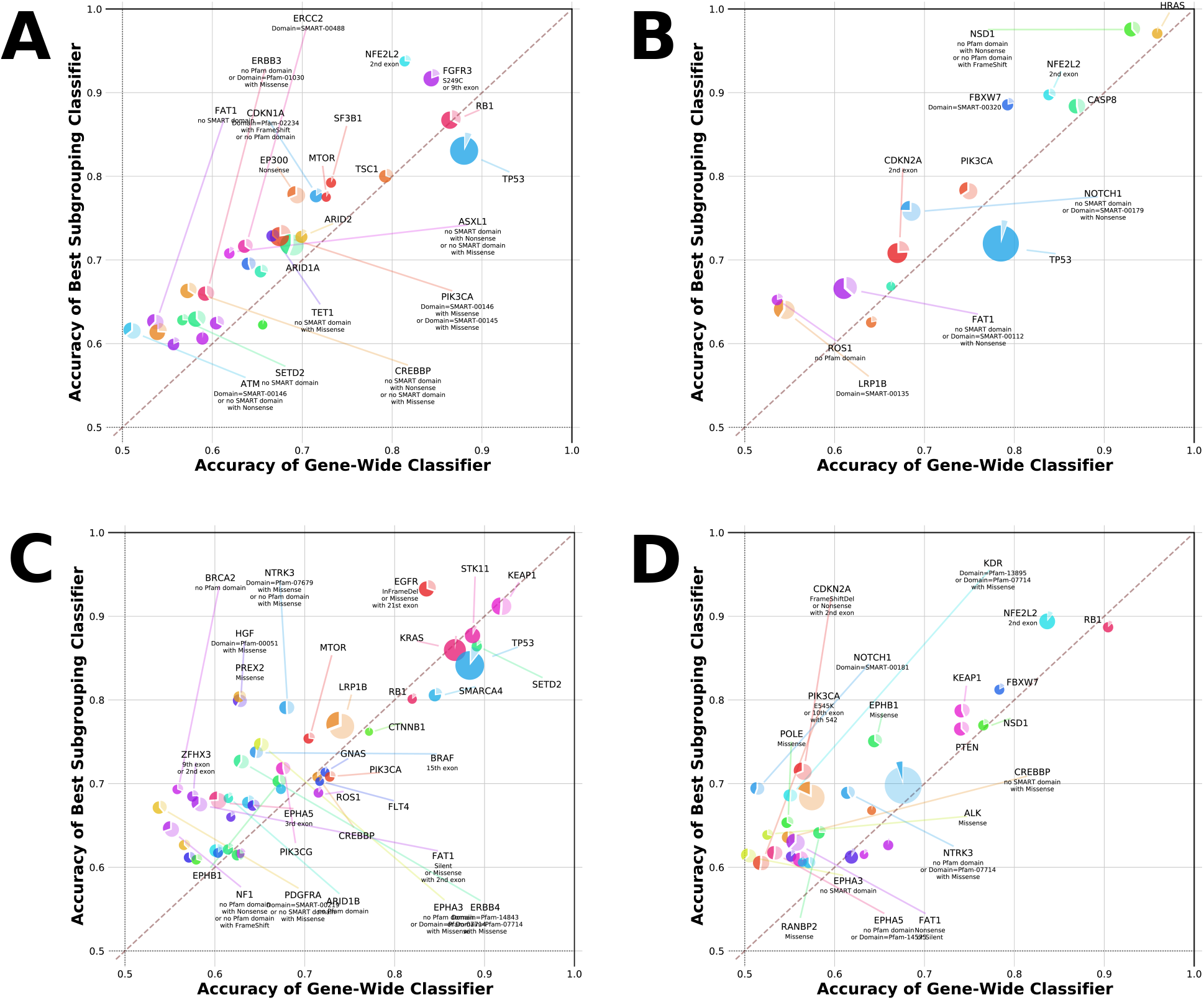
Divergent subgrouping behaviour is present in many cancer cohorts. We repeated the subgrouping enumeration and classification experiment to characterize alteration divergence in TCGA cohorts such as (**A**) BLCA, (**B**) HNSC(HPV-), (**C**) LUAD, and (**D**) LUSC.

**FIG. S13.**
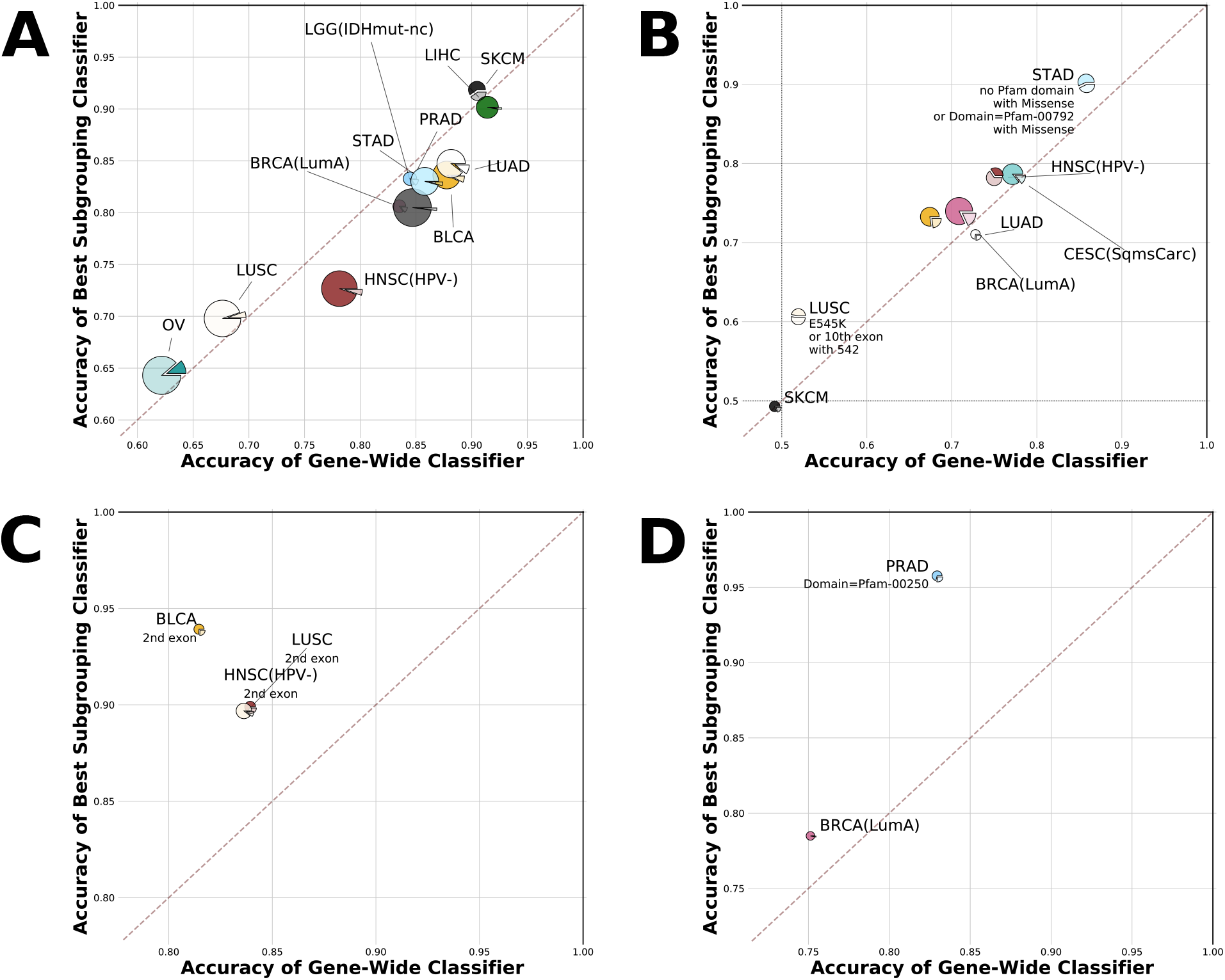
Comparing cancer gene perturbome characteristics across tumor contexts. The best subgroupings for (**A**) TP53, (**B**) PIK3CA, (**C**) NFE2L2, and (**D**) FOXA1 across the TCGA cohorts considered in this study. Each pie chart in a panel represents a cohort in which subgrouping mutation classifiers were trained and tested for the gene in question, with pie charts scaled and sliced according to the same schema as in Figure 1. The best found subgrouping of a gene within a cohort is listed wherever its down-sampled confidence score against the corresponding gene-wide classifier exceeded 0.8.

**FIG. S14.**
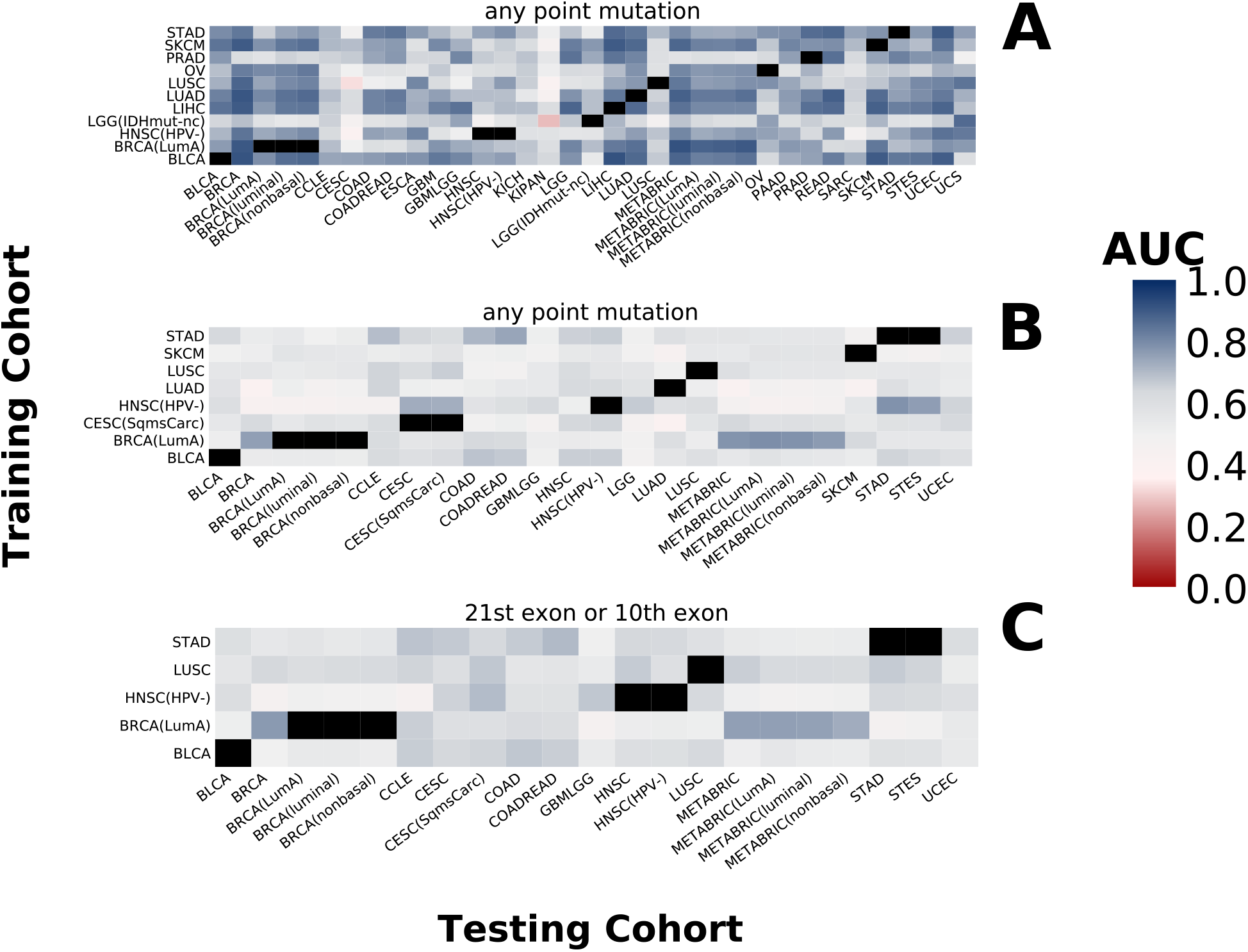
Transferring mutation signatures across disease contexts. Models trained to predict the presence of mutations and their subgroupings in each cohort were applied to every other cohort in which the corresponding mutation was also present. (**A**) The AUC performance of gene-wide TP53 classifiers according to the training cohort (x-axis) and the cohort they were transferred to (y-axis). (**B**) The AUC performance of gene-wide PIK3CA classifiers according to training cohort and transfer cohort as above. (**C**) Transfer AUC performance of the optimal PIK3CA subgrouping found by aggregating downsampled confidence scores between PIK3CA subgroupings and gene-wide tasks across all cohorts in which PIK3CA subgroupings were enumerated.

**FIG. S15.**
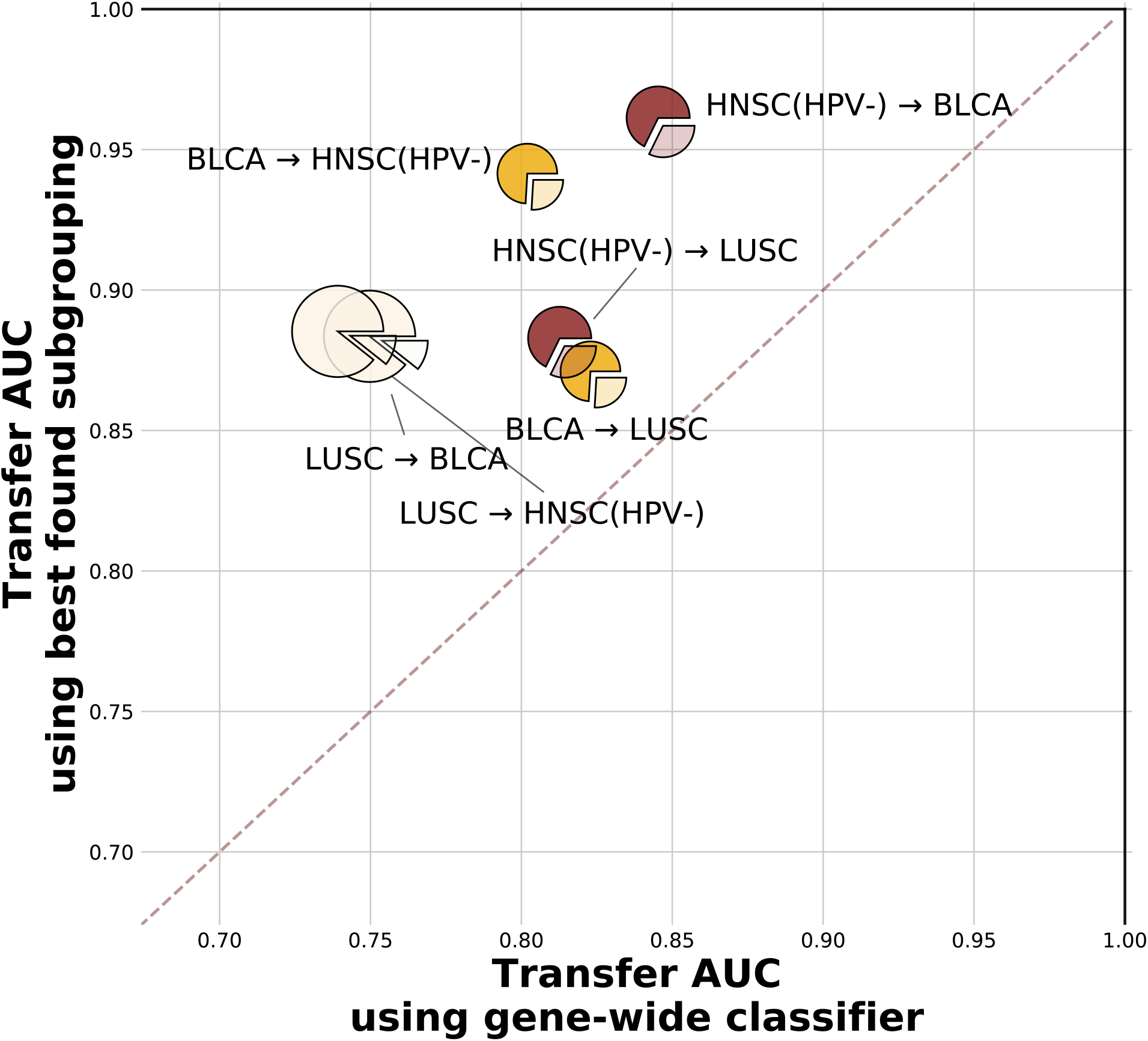
NFE2L2 subgrouping divergence is consistent across transfer contexts. Transfer performance of the optimal NFE2L2 subgrouping consisting of all mutations on the 2nd exon relative to the transfer performance of the NFE2L2 gene-wide task. Each pie chart corresponds to an instance of training the gene-wide and best found subgrouping classifiers in one TCGA cohort then asking them to make predictions in another TCGA cohort. Pie charts are sized according to the proportion of samples carrying any point mutation of NFE2L2 in the training cohort, with slices denoting the proportion of NFE2L2 mutants belonging to the optimal subgrouping in the training cohort.

**FIG. S16.**
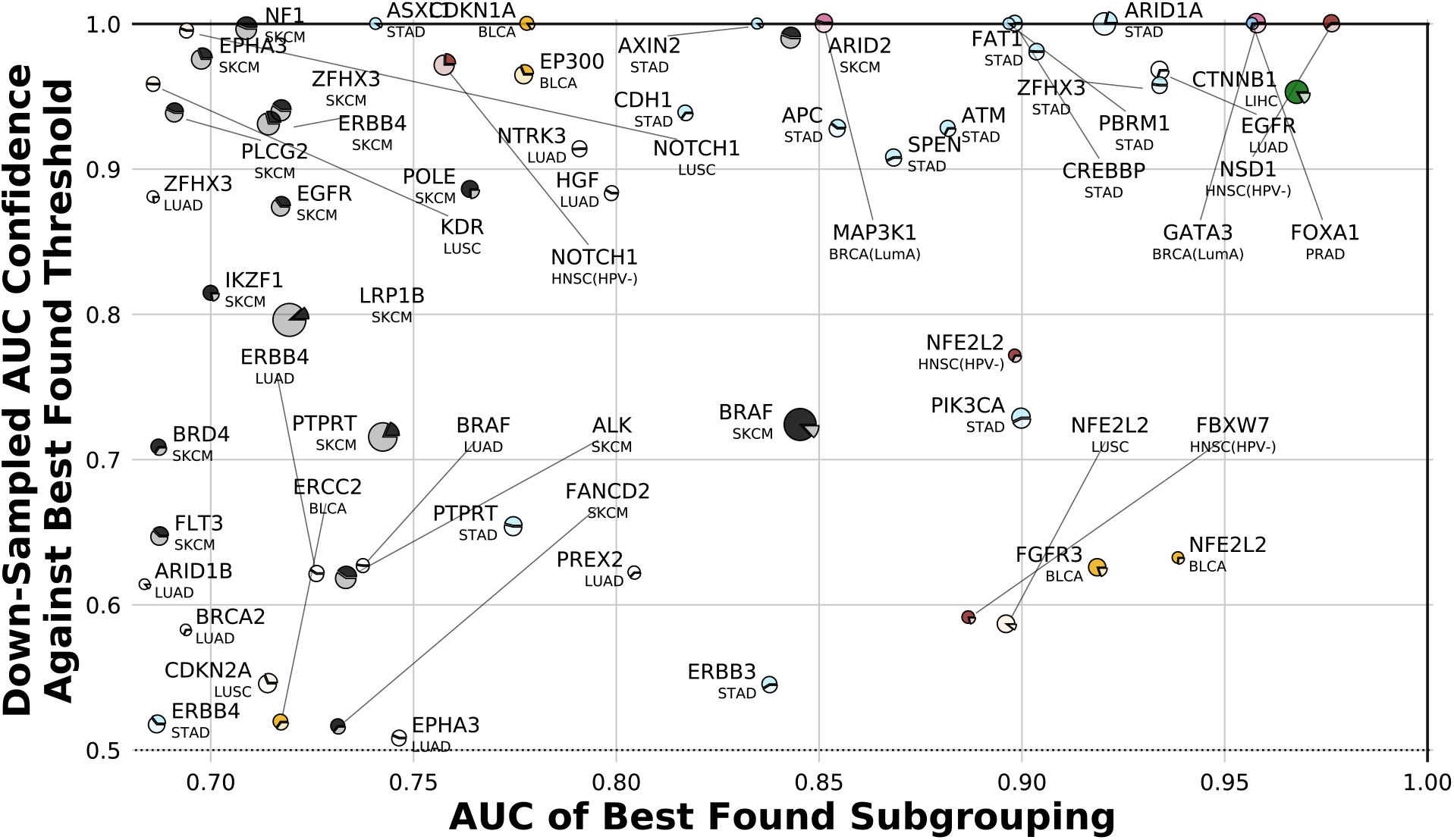
Cataloguing cancer genes where hierarchically-chosen subgroupings outperform subgroupings chosen using variant significance metrics. In each cancer cohort in which we looked for subgroupings we compared the performance of each gene’s optimal subgrouping’s expression classifier against that of the best threshold subgrouping chosen by considering other metrics of mutation significance (PolyPhen and SIFT). To quantify the significance of the difference between each pair of AUCs, we calculated the probability that a downsampled AUC for the best found subgrouping was higher than a downsampled AUC for the best found PolyPhen/SIFT threshold mutation subset. Results have been filtered to only include cases where the optimal subgrouping found using our mutation property hierarchies outperforms the gene-wide classifier (downsampled AUC confidence of at least 0.8 using the same approach).

**FIG. S17.**
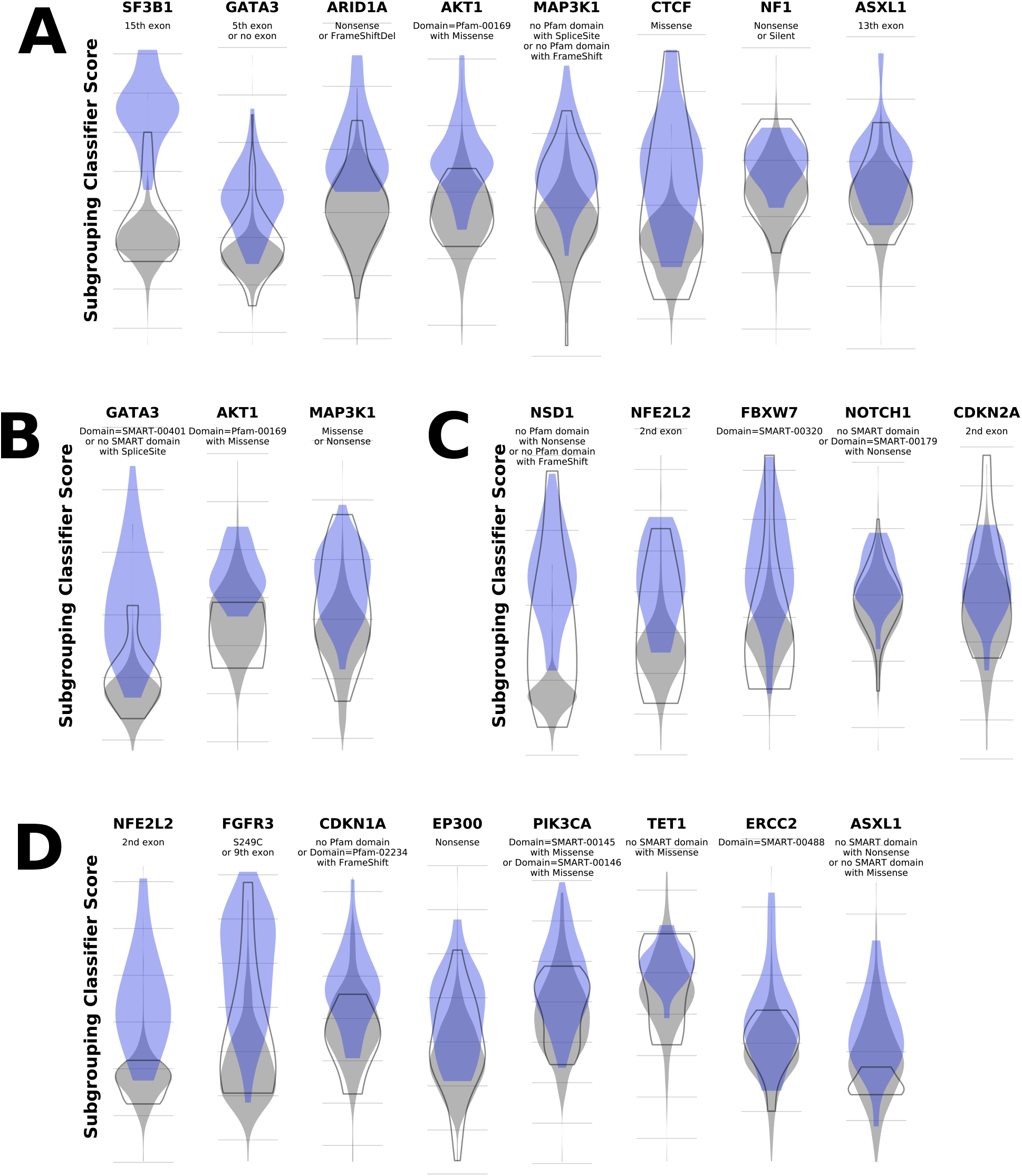
Subgrouping classifier scores reveal relationships between subgroupings and other mutations on the same gene. For genes with divergent subgroupings in (**A**) METABRIC(LumA), (**B**) TCGA-BRCA(LumA), (**C**) TCGA-HNSC(HPV-), and (**D**) TCGA-BLCA we considered the distributions of scores assigned by the best found subgrouping’s classifier to samples with the subgrouping’s mutations (blue violins), samples with point mutations on the same gene but not in the subgrouping (empty grey violins), and samples that are wild-type for point mutations on the gene (filled grey violins).

**FIG. S18.**
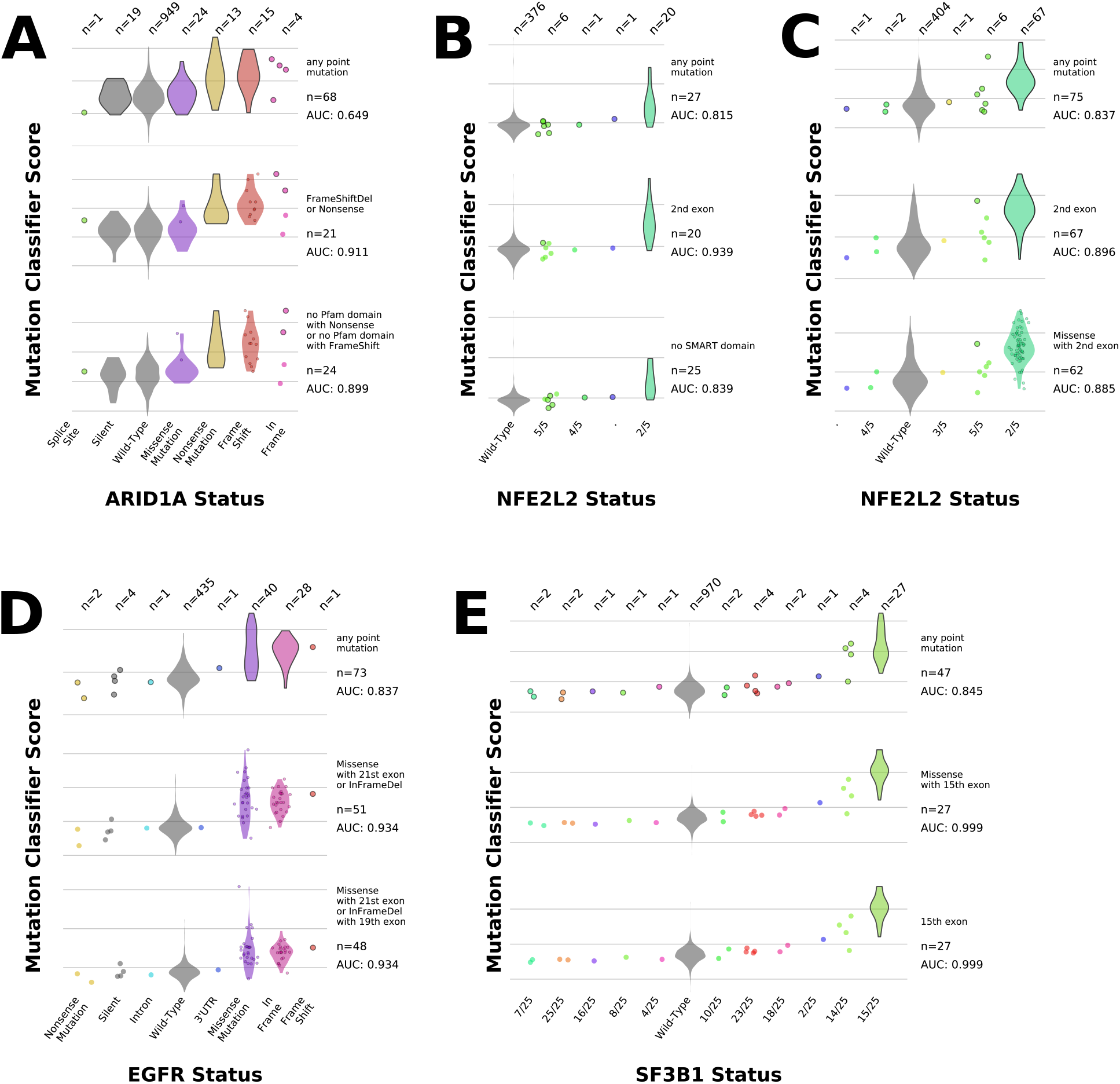
Subgrouping classifier scores reveal relationships between mutations within cancer genes. We dissected the scores returned by our mutation classifiers for mutations within (**A**) ARID1A in METABRIC(LumA), (**B**) NFE2L2 in TCGA-BLCA, (**C**) NFE2L2 in TCGA-LUSC, (**D**) EGFR in TCGA-LUAD, and (**E**) SF3B1 in METABRIC(LumA). Within each panel, rows correspond to classification tasks, with the top row showing scores for the gene-wide task and the remaining rows showing the best found subgroupings for the gene in question. Cohort samples are divided across the panel columns according to the type of mutation on the gene they carry, if any. Points and violins with a dark outline denote samples and populations of samples respectively that carried mutations the task had to predict; if a population contained mutated samples that were in the subgrouping as well as samples that were not in it then the samples in the subgrouping are plotted as points within the violin, which contains all samples in the population in every case.

**FIG. S19.**
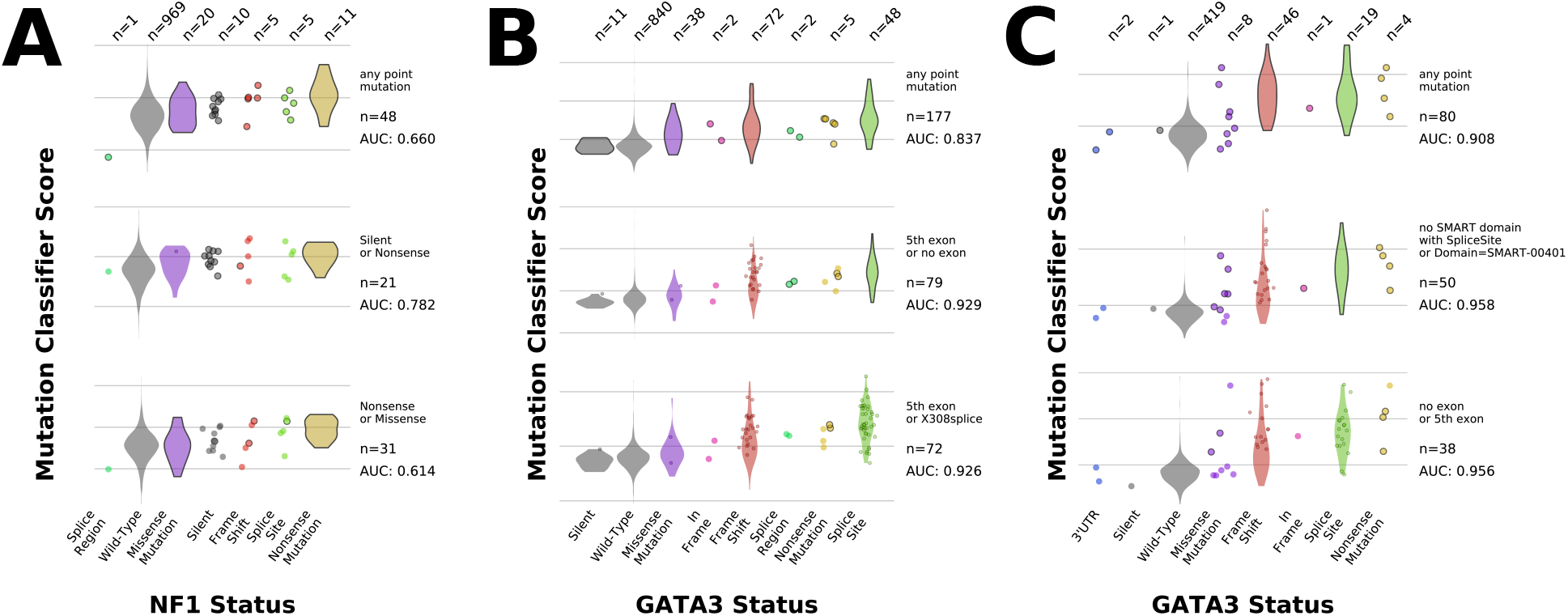
Subgrouping classifier scores reveal relationships between different classes of mutations within NF1 and GATA3. Scores returned by our mutation classifiers are plotted in the same style as in Figure S18 for mutations within (**A**) NF1 in METABRIC(LumA), (**B**) GATA3 in METABRIC(LumA), and (**C**) GATA3 in TCGA-BRCA(LumA).

