## Supplementary figures and images for "Systematic interrogation of mutation groupings reveals divergent downstream expression programs within key cancer genes"

### Graphical Abstract

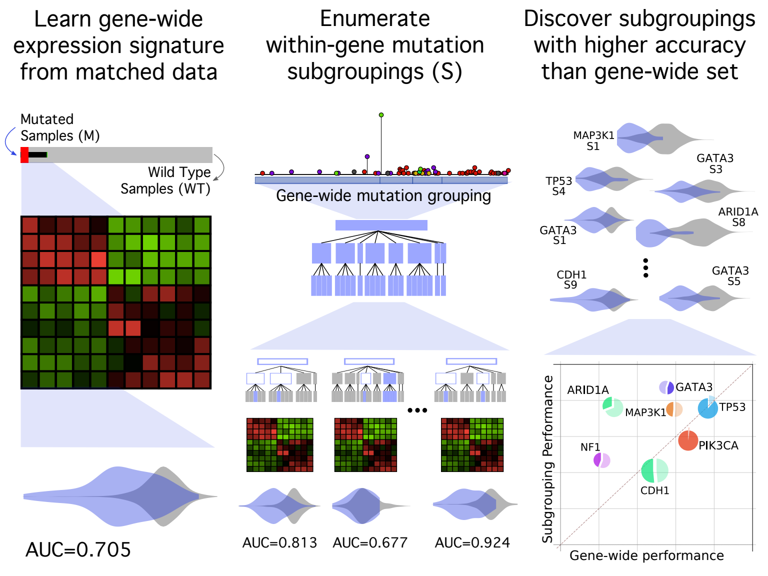
